# Higher-order epistasis within Pol II trigger loop haplotypes

**DOI:** 10.1101/2024.01.20.576280

**Authors:** Bingbing Duan, Chenxi Qiu, Steve W. Lockless, Sing-Hoi Sze, Craig D. Kaplan

## Abstract

RNA polymerase II (Pol II) has a highly conserved domain, the trigger loop (TL), that controls transcription fidelity and speed. We previously probed pairwise genetic interactions between residues within and surrounding the TL for the purpose of understand functional interactions between residues and to understand how individual mutants might alter TL function. We identified widespread incompatibility between TLs of different species when placed in the *Saccharomyces cerevisiae* Pol II context, indicating species-specific interactions between otherwise highly conserved TLs and its surroundings. These interactions represent epistasis between TL residues and the rest of Pol II. We sought to understand why certain TL sequences are incompatible with *S. cerevisiae* Pol II and to dissect the nature of genetic interactions within multiply substituted TLs as a window on higher order epistasis in this system. We identified both positive and negative higher-order residue interactions within example TL haplotypes. Intricate higher-order epistasis formed by TL residues was sometimes only apparent from analysis of intermediate genotypes, emphasizing complexity of epistatic interactions. Furthermore, we distinguished TL substitutions with distinct classes of epistatic patterns, suggesting specific TL residues that potentially influence TL evolution. Our examples of complex residue interactions suggest possible pathways for epistasis to facilitate Pol II evolution.

## INTRODUCTION

Functional interactions among amino acid residues within a protein can be detected by genetic interactions in the form of epistasis. Epistasis between residues can dmodulate the phenotypic effects of mutations. Specific substitutions can create or relax functional constraints on the identity of residues at other positions in a protein, and impact evolutionary trajectories (MANI *et al*. 2008; PHILLIPS 2008; STARR AND THORNTON 2016; DOMINGO *et al*. 2019; PARK *et al*. 2022; JOHNSON *et al*. 2023; METZGER *et al*. 2023). Epistasis is the modulation, positive or negative, of the effect of one mutant by the presence of other mutants. This phenotypic modulation can be detected by observing differences in the phenotype of a double mutant from that expected from the cumulative effects of the two corresponding single mutants, which is the baseline expectation in the absence of epistasis (PHILLIPS 1998; HILL *et al*. 2008; MANI *et al*. 2008; PHILLIPS 2008). Prior studies found many epistatic interactions at both lower-orders (i.e. among residue pairs) and higher-orders (among many residue combinations). Higher-order epistasis, emerging from particular residues, can reflect specific biological or physical interactions and imply crucial roles of these residues in proteins’ function and evolution (MELAMED *et al*. 2013; OLSON *et al*. 2014; BANK *et al*. 2015; POKUSAEVA *et al*. 2019; REDDY AND DESAI 2021; BAKERLEE *et al*. 2022; DING *et al*. 2022; LIN *et al*. 2022). In this work, we dissect complex higher-order epistasis by deep mutational scanning within an ultra-conserved and crucial active site domain, the trigger loop (TL), of RNA polymerase II (Pol II).

Pol II transcribes eukaryotic DNA into mRNA using an iterative nucleotide addition cycle (NAC) (BAR-NAHUM *et al*. 2005; WANG *et al*. 2006; MALINEN *et al*. 2012; DANGKULWANICH *et al*. 2013; KAPLAN 2013; KULDELL AND KAPLAN 2024). The TL is a critical domain for the NAC, and participates in all three steps: nucleotide selection, catalysis, and likely translocation by switching between different conformations (WANG *et al*. 2006; FOUQUEAU *et al*. 2013; BARNES *et al*. 2015). In each NAC, a matched substrate entering the active site induces or captures a conformational change of the TL from an open, catalysis-disfavoring state to a closed, catalysis-favoring state. Thus, the TL functions in kinetic selection to distinguish matched NTPs complementary to the DNA template from mismatched ones (WANG *et al*. 2006; HUANG *et al*. 2010; MALINEN *et al*. 2012; KAPLAN 2013; WANG *et al*. 2013; XU *et al*. 2014; BARNES *et al*. 2015). Full closure of the TL promotes catalysis (VASSYLYEV *et al*. 2007; WANG *et al*. 2013). Following catalysis, the TL transits from the closed to the open state, facilitating polymerase translocation for the subsequent NAC (SEIBOLD *et al*. 2010; DA *et al*. 2012; LIU *et al*. 2016) (**Fig. 1A**).

**Figure 1.**
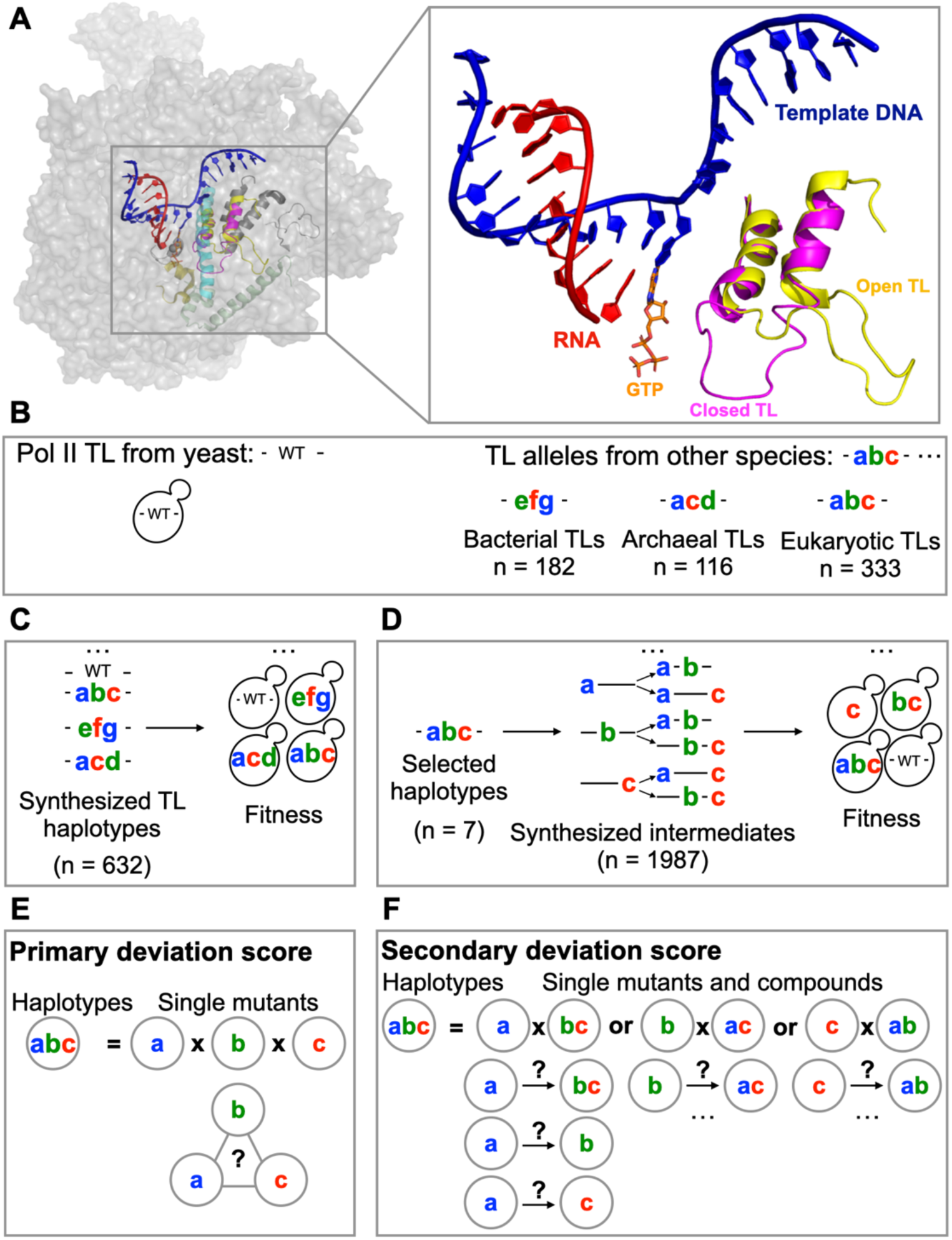
Systematic detection of TL-internal epistasis with natural TL alleles and intermediates. **A**. The Pol II TL is the key domain balancing transcription speed and fidelity in Pol II active site. Left panel: yeast Pol II structure (PDB: 5C4X). Right panel: The open, catalytic disfavoring (PDB: 5C4X) (BARNES *et al*. 2015) and closed, catalytic favoring (PDB: 2E2H) (WANG *et al*. 2006) conformations of TL in the active site. **B**. We selected 632 TL haplotypes representing TL alleles from bacterial, archaeal and the three conserved eukaryotic msRNAPs to detect TL-internal epistasis (Bacterial TLs n = 182, archaeal TLs n = 116, Pol I TLs n = 144, Pol II TLs n = 94, Pol III TLs n = 95). **C**. The selected TL alleles were synthesized and transformed into yeast Pol II to form chimeric Pol II enzymes. Yeast chimeric Pol II enzymes were phenotyped under selective conditions to detect growth defects, which are represented by fitness (see **Methods**). **D**. Seven Pol II TL alleles were selected to construct intermediate haplotypes representing all possible combinations of substitutions of the seven selected alleles. The intermediates were transformed into yeast to measure growth defects as in C. E-F. Analytical scheme of primary deviation score (**E**) and secondary deviation score (**F**). Details are in **Methods**.

Additional TL conformations have been associated with other RNA polymerase activities such as pausing and backtracking (WANG *et al*. 2009; ZHANG *et al*. 2010; CHEUNG AND CRAMER 2011; MOSAEI AND ZENKIN 2021). With these transitions, the mobile TL balances transcription fidelity and speed (WANG *et al*. 2006; KAPLAN *et al*. 2008; KIREEVA *et al*. 2008; SYDOW *et al*. 2009; KAPLAN 2010; YUZENKOVA *et al*. 2010; LARSON *et al*. 2012; UNARTA *et al*. 2023; KULDELL AND KAPLAN 2024).

How do residues within the TL interact with each other to ensure its proper transition? Our observations of distinct pairwise interactions suggest that TL function is maintained by residue interaction networks (DUAN *et al*. 2023). For example, previous biochemical and genetic studies found mutations in the TL can modify its folding or dynamics, causing changes in catalytic activity. These changes include increases to catalytic rate, leading to faster elongation than WT in vitro (gain of function, GOF) or decreases to catalytic rate, leading to slower elongation than WT in vitro (loss of function, LOF) (KAPLAN *et al*. 2008; KIREEVA *et al*. 2008; KAPLAN *et al*. 2012; BRABERG *et al*. 2013; NAYAK *et al*. 2013; WINDGASSEN *et al*. 2014; BARNES *et al*. 2015; QIU *et al*. 2016). Our prior work on TL double mutants, including combinations between or within GOF and LOF mutations, identified distinct types of pairwise interactions. These interactions resulted in fitness levels that were either better or worse than expected, based on the simple assumption that independently functioning substitutions would have cumulative effects. This indicates that mutations interact in ways that alter fitness beyond the additive effects. These interactions include suppression, commonly observed in certain GOF/LOF combinations, which leads to better than expected fitness. The suppressive interactions support a model where single mutants act independently within double mutants. In this model, the effects are balanced in double mutants that consist of mutations with opposite effects. However, in some GOF/LOF combinations, we observed a lack of suppression, known as sign epistasis, indicating the effect of one mutant is controlled by the identity of another residue. In contrast, in double mutants with similar phenotypes (GOF/GOF or LOF/LOF), the effects are enhanced to a level that is similar or even beyond the additivity. This latter situation is known as synthetic sickness or lethality. However, we also observed a lack of enhancement for some GOF/GOF or LOF/LOF combinations, referred to as epistasis, revealing dependency between residues (KAPLAN *et al*. 2012; QIU *et al*. 2016; QIU AND KAPLAN 2019; DUAN *et al*. 2023).

The TL is highly conserved across evolution, supporting the idea that identities of key residues are highly constrained (PALO *et al*. 2021). Given this conservation, it was surprising that analogous mutations in a highly conserved residue yielded opposite effects in the conserved yeast Pol I and Pol II enzymes (VIKTOROVSKAYA *et al*. 2013). Given the TL’s flexible and mobile character, along with its high conservation of sequence and function, it was an open question of if the TL is functionally self-contained, or evolutionary coupling through non-conserved residues is necessary to maintain its ability to achieve different conformations. Our recent studies identified widespread incompatibility between trigger loops of different species or enzymes placed into the yeast Pol II context, supporting the idea of an epistasis network from residues outside the TL is prominent in contributing to TL function and creating different contingencies for its evolution (DUAN *et al*. 2023). We wished to understand the drivers of this incompatibility between conserved TLs and use diverse TL haplotypes to probe the complexity of internal TL interactions.

Recent studies suggest that the primary contributor of epistasis to protein function is through pairwise residue interactions. However, higher order epistasis is complicated and can be difficult to detect (JOHNSON *et al*. 2023; METZGER *et al*. 2023). We reasoned that the Pol II TL might be an excellent system to detect higher order epistasis using evolutionary haplotypes, notwithstanding constraints on function from the rest of the enzyme complex. The Pol II TL is flexible, required to support distinct conformational states, and functionally plastic, as mutants can both increase or decrease catalysis. Notably, known suppressor relationships between these two types of mutants highlight the complexity of residue interactions within the TL. We applied our TL deep mutational scanning system to detect higher order epistasis by comprehensive dissection of a set of evolutionarily derived TL sequences (haplotypes). We identified intra-TL interactions of evolutionarily observed substitutions (TL-internal epistasis) and dissected specific examples of complex interactions that would not be evident from simple analyses only considering single mutants or complete haplotypes. Our experiments suggest specific paths for TL evolution in the context of the complete enzyme may likely go through mild gain-of-function residues that allow additional changes to be tolerated.

## RESULTS

### Systematic detection of epistatic interactions within TL haplotypes

We utilized our previously developed deep mutational scanning-based phenotyping platform for Pol II mutants in *Saccharomyces cerevisiae* (QIU *et al*. 2016; DUAN *et al*. 2023) to analyze TL-internal higher-order epistasis. The analyzed TL haplotypes included 632 natural TL variants from bacterial and archaeal multi-subunit RNA polymerases (msRNAPs), and eukaryotic Pol I, Pol II and Pol III (**Fig. 1B-C**). An additional 1987 TL alleles were included, representing all possible intermediate substitution combinations for seven selected natural Pol II TL variants (**Fig. 1D**). The seven TL variants were selected because they contain specific amino acids which exhibit phenotypes, either GOF or LOF, when introduced individually in yeast Pol II. The functional consequences of these haplotypes were measured by a set of growth phenotypes that are predictive of biochemical functions (see **Methods**)(KAPLAN *et al*. 2012; QIU *et al*. 2016; QIU AND KAPLAN 2019; DUAN *et al*. 2023). The low coefficient of variation values observed in all libraries indicates high reproducibility for our measurements (**Fig. S1A**). Our system allows us to profile these haplotypes and assess epistasis among multiple mutations in a highly parallel fashion.

To detect epistasis, we first fitted a log additive model for the fitness scores of individual mutations, based on the assumption that se mutations acting independently will have log additive effects on fitness(HILL *et al*. 2008; MANI *et al*. 2008; PHILLIPS 2008; LIN *et al*. 2022; DUAN *et al*. 2023). We then calculated deviations from the expected fitness scores by comparing the observed fitness of haplotypes to these expectations (**Fig. S1B**). Any deviation from the log additive model is evidence for residue interactions (epistasis). Deviation from expectation could be in the form of mutual suppression (double mutant more fit than either single mutant), basic epistasis (double mutant no worse than one of the single mutants), or synthetic sickness (greater than log additive defects in the double mutant). The deviation values, termed “deviation scores”, were calculated to detect intra-TL genetic interactions within haplotypes (see **Methods**).

Deviation scores were calculated in two ways. First, we determined a “primary deviation” score to represent total epistasis within a haplotype. This score quantifies the deviation between the observed fitness of the haplotype to its expected fitness, which is calculated by summing the fitness scores of the individual mutants involved (**Fig. 1E and Fig. S1B**). Second, to determine which residues the epistasis emerged from, we separated the haplotypes into combinations of single substitutions and the compound mutants formed by the remaining substitutions. We therefore calculated a secondary deviation score. This score is determined by comparing the observed fitness score of the haplotype to the sum of two components: the fitness score of a single substitution and the fitness score of the compound formed by the remaining substitutions (**Fig. 1F and Fig. S1B**). The magnitude of the secondary deviation score indicates the magnitude of epistatic effects for specific substitutions when they are introduced to the compound mutant. Moreover, analyzing the secondary deviation scores of a specific substitution across various compound mutants (backgrounds) enables us to assess the breadth of its epistatic effects, which may reveal its potential to impact TL evolution.

### Drivers of incompatibility for TL variants placed in the yeast Pol II context

We investigated the widespread incompatibility of TL haplotypes from various msRNAPs when introduced into yeast Pol II (VIKTOROVSKAYA *et al*. 2013; QIU *et al*. 2016; DUAN *et al*. 2023). To probe the incompatibility in detail, we grouped haplotypes based on their evolutionary source (bacteria, archaea and eukaryotic Pol I, Pol II and Pol III). We then illustrate the fitness and deviation scores in heatmaps, including the fitness of all constituent single substitutions as measured in yeast Pol II (QIU *et al*. 2016; DUAN *et al*. 2023), the expected fitness of the haplotypes calculated using the log additive model, the observed fitness of the haplotypes measured from our deep mutational scanning, and the primary deviation score derived from comparing of the observed to the expected fitness (**Fig. 2A-E**). We found that the incompatibility of bacterial TLs in yeast could be explained by three substitutions that are individually lethal in yeast (**Fig. 2A**). In contrast, only about half of the archaeal TLs had lethality that could be explained by individually lethal substitutions. The other half appeared to be lethal due to the straightforward additivity of single mutant fitness, although a subset exhibited negative epistasis among the constituent substitutions (**Fig. 2B**). Eukaryotic Pol I, Pol III and Pol II haplotypes exhibited fewer individually lethal single substitutions than those in bacteria and archaea. However, they displayed more instances of both positive and negative intra-TL interactions (**Fig. 2C-G**), likely reflecting the closer evolutionary distance of these eukaryotic Pols to yeast Pol II. Interestingly, a few single substitutions from other Pol II TL haplotypes caused lethality when introduced into yeast Pol II context (**Fig. 2D**). We examined whether the fitness of Pol II haplotypes correlates with the evolutionary distance between the genes where the TL haplotypes originate and *RPB1*, where the WT TL is derived. However, we did not see a significant correlation (**Fig. S2**). This could be because the TL haplotypes we picked were chosen based on the diversity of single substitutions, rather than an unbiased selection across evolutionary distances from *S. cerevisiae RPB1*. Therefore, we have some selection bias in our haplotypes. In summary, our analyses indicate that, despite the high conservation across Pol II enzymes, specific and important functional constraints have evolved among close homologs, emphasizing the significance of epistasis between the TL and its Pol II context.

**Figure 2.**
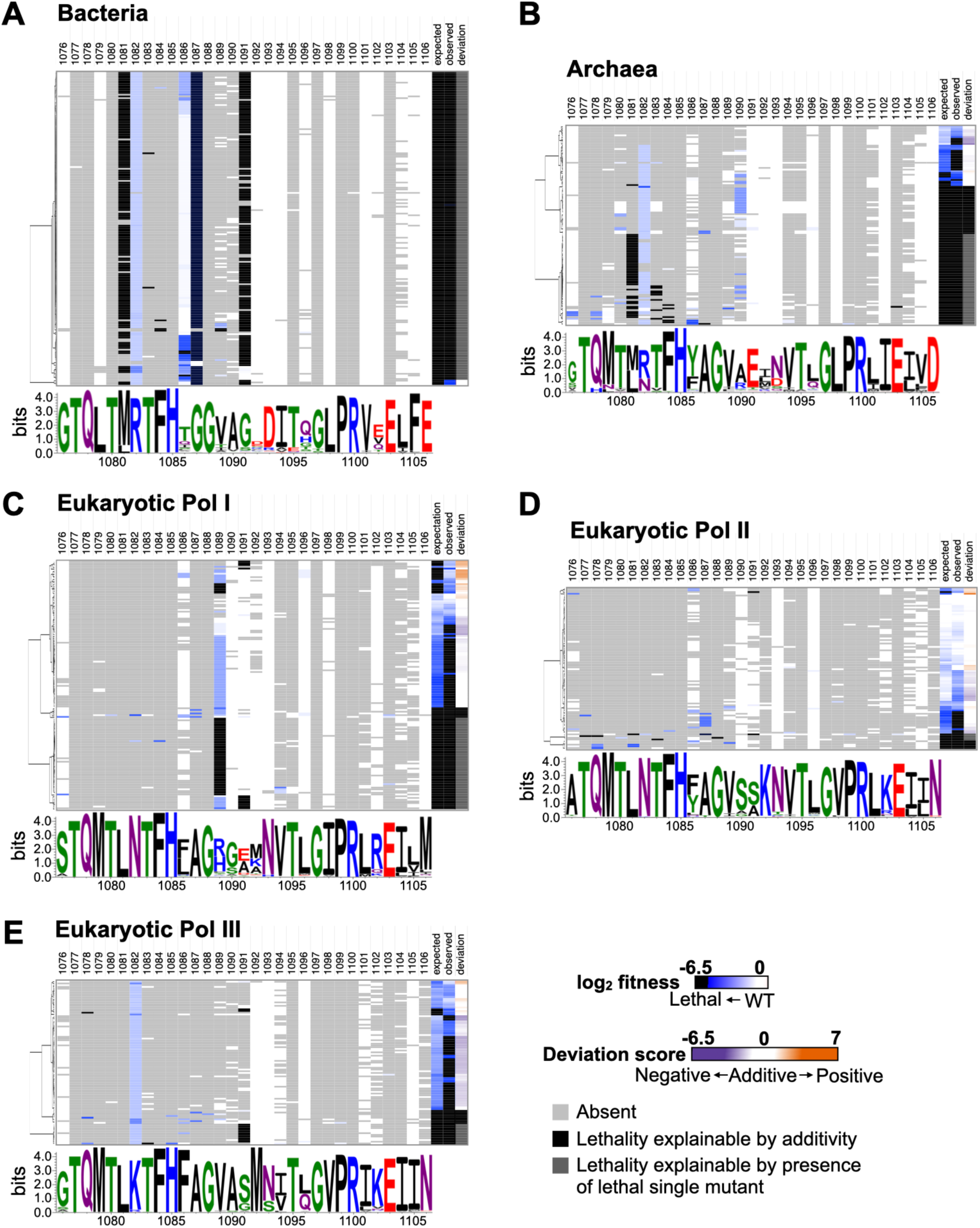
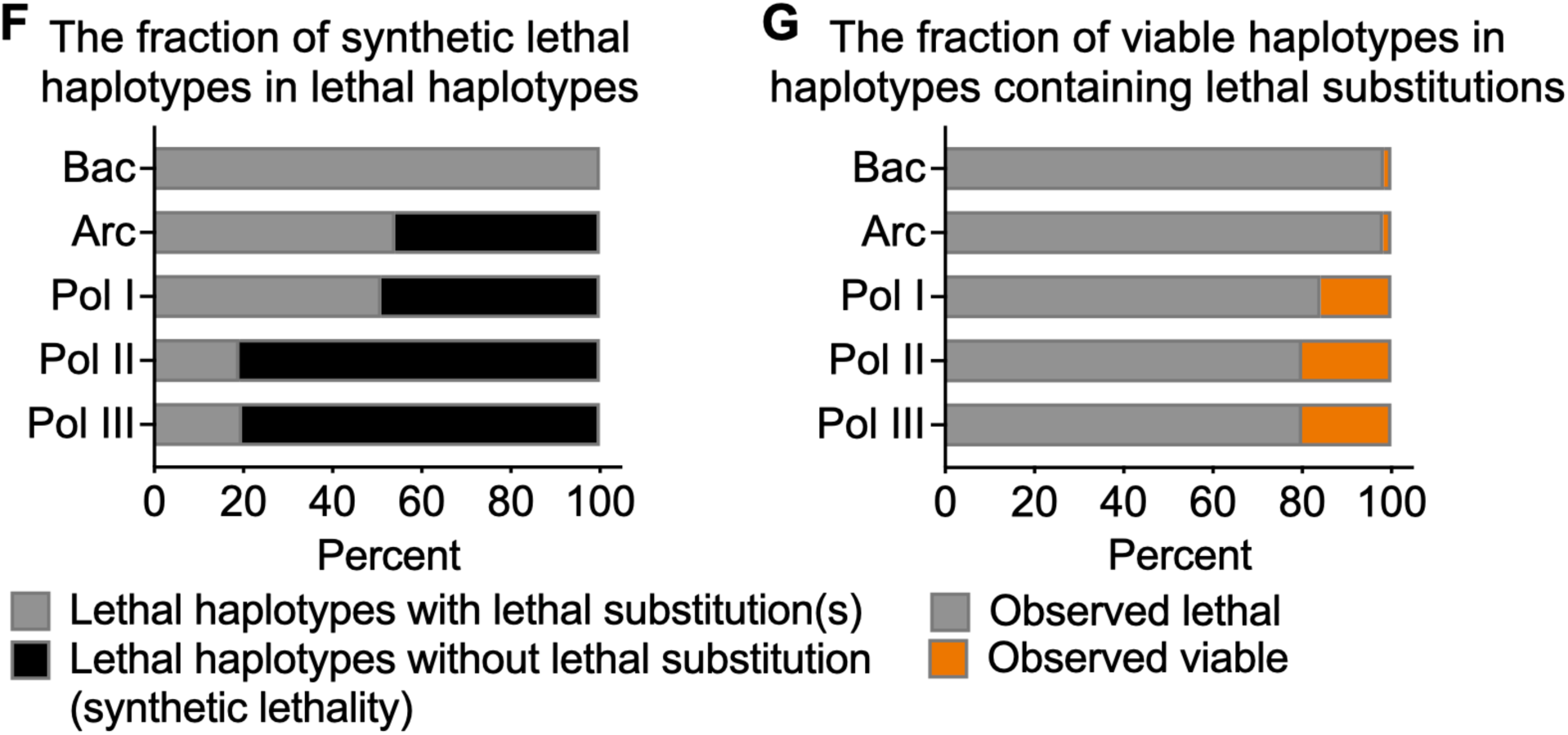
Greater TL-internal epistasis is observed with closer evolutionary distance to eukaryotic Pol II. **A-E**. Fitness and primary deviation score heatmaps of TL haplotypes from five msRNAP evolutionary groups. The *x*-axis of each heatmap is the 31 residue positions of the Pol II TL (1076-1106). The *y*-axis of each heatmap is the TL haplotypes belonging to each group clustered by hierarchical clustering with Euclidean distance. Each row represents one haplotype with several single substitutions. Light grey blocks in each row represent the residue from the haplotype at the position is the same with yeast Pol II TL residue, in other words, no substitution at the position. Colored blocks represent different residues in the haplotype compared with yeast Pol II TL (substitutions). The color of the block represents growth fitness of the single substitution in the yeast Pol II TL background. Expected fitness of the haplotypes were calculated from the log additive model for individual substitutions. Observed fitness was measured in the screening experiments, and deviation scores were calculated by comparing the observed and expected scores (shown at the right end of each row). Sequence logos were generated with multiple sequence alignment (MSA) of the five groups individually in Weblogo 3.7.12. The labeled numbers of the sequence logo represent yeast Pol II TL residue position (1076-1106). Bacteria n=465. Archaea n=426. Pol I n=605. Pol II n=405. Pol III n=444. **F**. Lethal haplotypes can contain three distinct types of lethality: one attributed to synthetic lethal interactions among substitutions, representing negative interactions that result in lethality beyond expectation, additive lethality expected from combination of individual substitutions, and that arising directly from the presence of an individually lethal single substitution. The calculated ratio specifically reflects the proportion of lethality due to negative interactions and additive lethality within all types of lethal haplotypes. **G**. The ratio of viable haplotypes in haplotypes containing lethal substitutions, representing the ratio of positive interactions (suppression) in TL haplotypes of each group. Haplotypes containing lethal substitutions are expected to be lethal based on the additive model. If haplotypes with lethal substitutions are observed to be viable, it suggests other substitutions suppress the lethal substitution in the haplotypes, implying positive epistasis (suppression). Approximately 20% of examined eukaryotic TL haplotypes containing individual lethal substitutions are viable, whereas only roughly 2% of bacterial and archaeal TL haplotypes are viable, suggesting positive interactions in eukaryotic TLs can buffer negative effects in more closely related TL sequences.

We then asked if functional coupling of residue identities within the TL across evolution might correlate with positive interactions (**Fig S3A-C**). This analysis tests the hypothesis that specific residues working together might be reflected by their coappearance in the TL, and if it might be observable independently from their coevolved greater enzymatic contexts. Statistical coupling analysis, which uses statistical inference to identify functional interactions by analyzing residue identities that correlate across evolution, served as our method (RUSS *et al*. 2005; SOCOLICH *et al*. 2005). Using a multiple sequence alignment of 362 eukaryotic TL sequences (natural TL variants), We identified 11 residues that exhibited coupling within the TL domain (**Fig. S3A**).

To further explore the role of TL-internal statistical coupling in its function, particularly when removed from their natural context, we designed two libraries as previously described (RUSS *et al*. 2005; SOCOLICH *et al*. 2005). The first library consisted of scrambled haplotypes that preserved TL-internal residue couplings and conservation as derived from natural TL variants (“Monte Carlo” scrambling library) (**Fig. S3C**). The second library maintained only conservation but not any TL-internal residue coupling (“Random scramble” library) (See **Methods**) (**Fig. S3B**). Both libraries satisfy the design parameters as indicated by residue coupling heatmaps (**Fig S3A-C**). If TL-internal coupling plays a significant role in supporting TL function, we would predict that haplotypes in the Monte Carlo scramble library to exhibit higher fitness on average than those in Random scramble library. However, the observed difference in fitness distributions between the two libraries was subtle and statistically insignificant (P-value = 0.3013). This result suggests that without the epistasis between TL alleles and their coevolved natural contexts, the TL-internal epistasis is weak or undetectable.

### Epistasis within TL haplotypes can be attributed to specific residues

We next determined the residues from which epistatic effects in TL haplotypes emerge. We selected seven Pol II TL variants, each containing 7∼9 substitutions relative to the *S. cerevisiae* Pol II TL, based on their diversity. We constructed 2169 combinations of all substitutions from the seven variants. Some of the substitutions are shared between haplotypes, leading to repeated combinations. We kept all unique haplotypes (n=1987) to ensure an equal initial allele frequency for all combinations. These seven variants exhibited varying degrees of compatibility, as determined by their fitness when replacing the yeast Pol II TL (**Fig. 3A-H)**. Four out of seven TL variants were incompatible in the yeast Pol II context for distinct reasons. The TL variant from *G. luxurians* was lethal, presumably because it encodes G1088S, which causes lethality on its own and was similarly deleterious in all intermediate genotypes (**Fig. 3A, C**). In contrast, the lethalities of TL variants from *B. anathema* and *R. solani* could not be explained by any individual lethal substitution.

**Figure 3.**
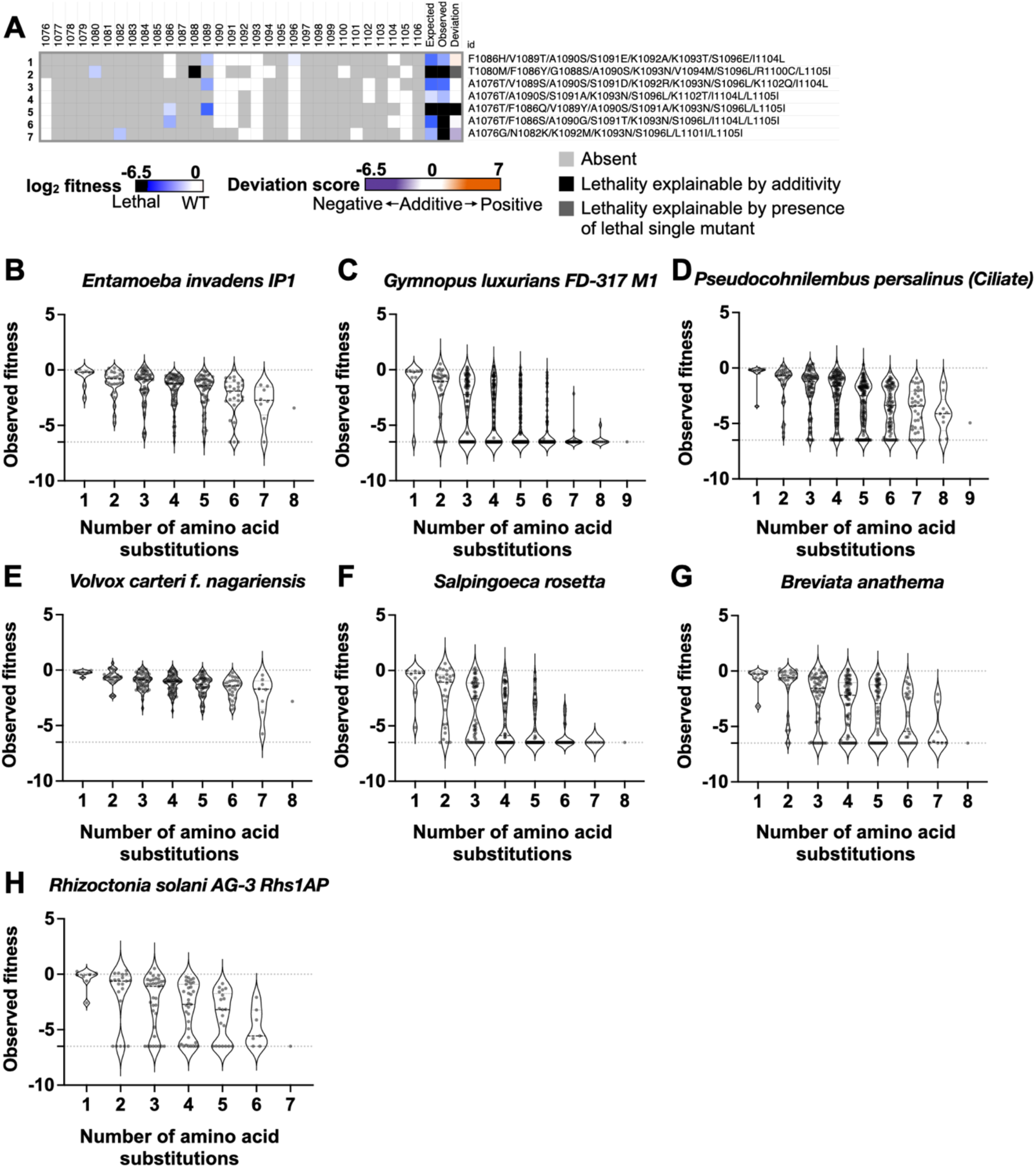
Phenotypes fluctuate with changes in substitution composition within haplotype groups. **A**. Fitness of constituent single substitutions, and expected and observed fitness of the haplotypes, and deviation score are shown in heatmap for the selected seven haplotypes. The deviations shown in the heatmap are the primary deviation scores calculated by comparing observed fitness to expected fitness. **B-H**. Distribution of growth fitness across all intermediate substitution combinations categorized by number of amino acid substitutions for each selected TLs. Each spot on the graph represents a haplotype. The *y*-axis indicates the fitness of each haplotype. The *x*-axis indicates the number of amino acid substitutions within these haplotypes. For example, consider a point in **B**, which represents a haplotype from the substitution combination in the TL of *E. invadens IP1*. If the point is positioned at x=3, showing it contains three substitutions, and at y=-0.5, indicating a fitness value of -0.5 for this haplotype.

They were predicted to be viable by the summation of the fitness scores of individual substitutions. This suggests that the observed lethality in these two haplotypes resulted from negative interactions among residues, indicating lethality beyond simple additivity (**Fig. 3A, G, H**). The *S. rosetta* TL, however, was both expected and observed to be lethal, suggesting that residue additivity causes lethality (**Fig. 3A).** Yet, it’s worth noting that some intermediate combinations of *S. rosetta* exhibited fitness in the lethal range **(Fig. 3F**) but were worse than predicted from the log additive model, suggesting negative interactions. Interestingly, the other three variants, where all individual substitutions were viable and combinations between them were predicted to be viable based on the log additive model, some intermediate substitution combinations fell into the severely sick or lethal range (**Fig. 3B, D, E and S4**). This implies negative interactions among the substitutions. However, the lethality of these intermediate genotypes was suppressed by additional substitutions, resulting in the complete, viable haplotypes and indicating positive interactions (**Fig. 3B, D, E**). The fluctuations of the fitness with various residue combinations suggests complexity and the potential for higher order epistasis within these TL haplotypes.

To identify which specific residues contributed to the observed epistasis, we assessed deviations from expected fitness when adding each substitution to intermediate combinations derived from the original haplotypes. This analysis enabled comparisons of mutant effects across multiple related backgrounds (**Fig. 1D, F and Fig. S1**). We then visualized the fitness and epistatic effects of each substitution using epistasis heatmaps (**Fig. 4 and S5**). For example, the *E. invadens IP1* TL haplotype exhibited better fitness than predicted by the log additive model (**Fig. 4A**), suggesting positive interaction within the eight substitutions in the haplotype. Among these substitutions, two have opposing biochemical phenotypes determined from library screening of single mutants: the LOF V1089T and the GOF S1096E. Notably, these two substitutions also displayed the lowest growth fitnesses among the eight *E. invadens IP1* TL substitutions (**Fig. 4A-B**). The characteristics of these alleles suggest they are likely candidates for strong epistatic effects. In support of this, S1096E consistently showed positive secondary deviation scores in most intermediate combinations, suggesting its broad effects. Conversely, V1089T primarily showed positive effects in combinations that contained S1096E (**Fig. 4B**), suggesting that V1089T’s positive effect is dependent on the presence of S1096E. This dependency is likely due to a suppression effect arising from the combination of GOF and LOF substitutions. Interestingly, S1091E, which lacks strong growth fitness defects and obvious phenotypes as a single substitution, showed strong positive or negative epistatic effects in most backgrounds (**Fig. 4B**). These results suggest that these three substitutions are key drivers of intra-TL epistasis within *E. invadens IP1* TL haplotype.

**Figure 4.**
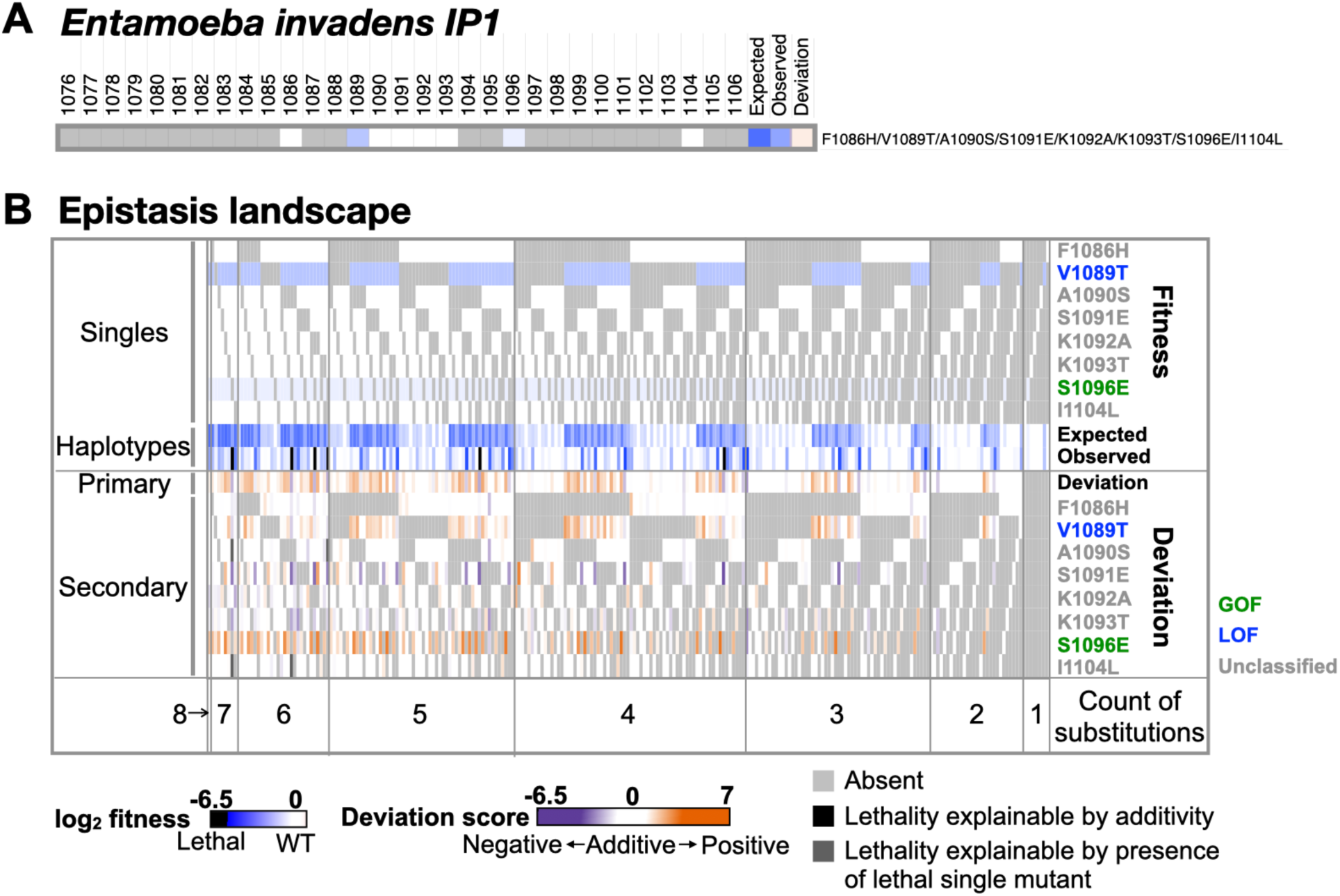
The epistasis landscape provides a comprehensive view of primary and secondary deviation scores, emphasizing substitutions with notable epistatic effects. **A**. Fitness of eight *Entamoeba invadens IP1* TL single substitutions in the yeast Pol II background, and the expected and observed fitness and the primary deviation score are shown in the heatmap. **B**. The epistasis landscape of *E. invadens IP1* TL substitutions. The heatmap illustrates the fitness and epistasis of all unique intermediate haplotypes coming from combinations of eight substitutions. Intermediate haplotypes are grouped by number of substitutions from 1 to 8. The fitness values are displayed in the upper panel and the epistasis, represented by primary and secondary deviation scores, is displayed in the lower panel. The colors of substitution names indicates their phenotype profiles as single mutants, GOF is in green, LOF is in blue, unclassified is in grey.

To visualize these effects, we quantified how individual substitutions influence epistasis within the haplotype. This was done by analyzing correlations between the primary deviation score, which reflects the overall epistasis, and the secondary deviation scores, which detail the specific epistatic effects of each substitution on different haplotype backgrounds. Substitutions demonstrating strong correlation (higher linear regression R^2^) are identified as the main drivers of epistatic effects observed in the entire haplotype. In contrast, substitutions with low correlation (smaller R^2^) are considered to have limited or negligible contributions. To illustrate this concept, we created a simplified simulation (**Fig. S5**). We constructed a TL haplotype comprising five substitutions: a, b, c, d, and e. In this scenario, the single mutant “a” is GOF, “b” and “c” are LOF, whereas “d” and “e” have no effects. We set the suppression interactions in GOF+ LOF combinations (a/b and a/c), and synthetic sick interactions in the LOF+ LOF combination b/c (**Fig. S5A**). We calculated both primary and secondary deviation scores for the haplotype and all intermediate combinations. Notably, the GOF mutant “a” exhibited positive secondary deviation scores in most intermediates (**Fig. S5B**). The secondary deviation scores of three single mutants with phenotypes showed strong correlation (R^2^) with the primary deviation scores (**Fig. S5C**), showing that they are the key contributors to the observed epistatic interactions within the haplotype. Further comparisons between the secondary deviation scores of “a” versus “b” and “c” revealed positive secondary deviation scores of “b” and “c” in most cases when “a” is present (**Fig. S5D**), suggesting potential suppression of “a” by “b” and “c”.

Similarly, the secondary deviation scores of “a” are consistently positive with changes in “b” (**Fig. S5E**) and “c” (**Fig. S5F**), indicating “a” suppresses “b” and “c”. These observations confirmed the mutual suppression in GOF and LOF combinations. Conversely, LOF substitutions “b” and “c” showed correlated secondary deviation scores, showing their synthetic sick interactions. These results demonstrate the behaviors that might be apparent for mutant types in this form of analysis.

To detect the key contributors to epistatic interactions within the *E. invadens IP1* TL haplotype, we determined correlations between primary and secondary deviation scores. Consistent with the epistasis landscape (**Fig. 4B**), the secondary deviation scores of S1096E, V1089T, and S1091E have strong correlations (R^2^ > 0.5) with the primary deviation score (**Fig. 5A**), suggesting their substantial contributions to epistasis within the haplotype. In all seven selected TL variants, one to three substitutions were identified that notably affect haplotype epistasis (**Fig 4B**, **5A, and Fig. S6-S7**), suggesting that epistasis within haplotypes is attributable to specific subsets of substitutions.

**Figure 5.**
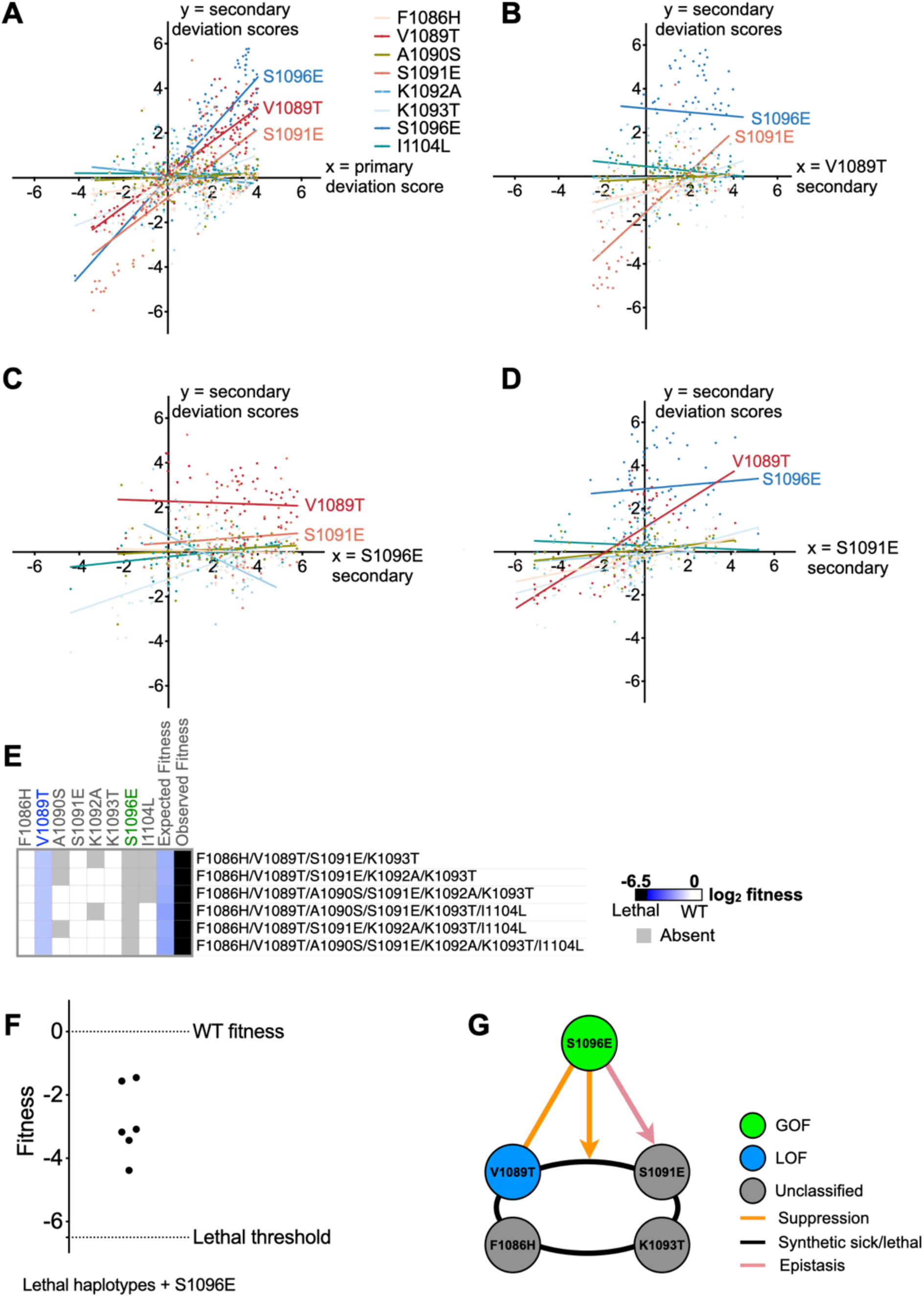
Correlations between deviation scores reflect specific residue interactions in *E. invadens IP1* TL substitutions. **A**. Correlations between secondary deviation scores of all eight substitutions (*y*-axis) and the primary deviation score (*x*-axis). Linear regression was applied to each comparison of secondary deviation scores against primary deviation scores to check the correlation. Substitutions with an R^2^ value exceeding 0.5 are annotated on the *x*-*y* plot, indicating their substantial impact on primary epistasis of the haplotypes. **B-D**. Correlations between secondary deviations of the other seven substitutions (*y*-axis) vs V1089T (**B**), S1096E (**C**), S1091E (**D**) on the *x*-axes respectively. **E**. The fitness landscape of intermediate combinations with fitnesses in the ultra-sick/lethal range. Their observed fitness levels are in the lethal range as indicated with black blocks in the heatmap while the expected fitness scores calculated from the additive model is in viable range (light blue blocks), indicating potential negative interactions. The fitness scores of the constituent single mutants within lethal haplotypes are shown in the heatmap. F1086H, V1089T, S1091E, and K1093T are present in each lethal haplotype while S1096E is absent. Names of substitutions are colored based on their phenotype profiles as single mutants. GOF: green. LOF: blue. No obvious phenotype: grey. **F**. The fitness of all ultra-sick to lethal haplotypes with S1096E incorporated is no longer in the ultra-sick/lethal range. **G**. Scheme of specific residue interactions within substitutions of *E. invadens IP1* TL.

### The epistasis within TL haplotypes can reflect intricate residue interactions

We dissected the interactions formed by the three substitutions S1096E, V1089T, and S1091E in the *E. invadens IP1* TL haplotype, revealing an intricate, higher-order epistasis network (**Fig. 5A-G**).

First, we confirmed the mutual suppression between the GOF S1096E and the LOF V1089T. S1096E consistently showed positive epistatic effects in haplotypes containing V1089T **(Fig. 5B**). V1089T exhibited a smaller but similar positive effect on haplotypes containing S1096E (**Fig. 5C**), showing mutual suppression between them. These observations are consistent with the predicted suppression between GOF and LOF mutants, as illustrated by the positive values for their combinations shown in **Fig. 4B**. Additionally, we observed negative interactions involving S1091E when combined with F1086H/V1089T/K1093T, in the absence of S1096E. This was indicated by low secondary deviation scores for S1091E when S1096E was included (**Fig. 5C**). However, S1096E exhibited consistent high scores with S1091E included (**Fig. 5D**). These observations are interpreted as S1096E being epistatic to S1091E. We further identified negative interactions in six intermediate combinations containing S1091E, as suggested by their fitness in the ultra-sick or lethal range beyond what the log additive model predicts (**Fig. 5E**).

Strikingly, F1086H, V1089T, K1093T were present in all these combinations, pointing to negative synergistic interaction (synthetic sickness or lethality) between the four substitutions. Conversely, S1096E was absent in all these combinations (**Fig. 5E**), implying that its presence suppresses the negative interaction. This suppression was further supported when the inclusion of S1096E in these combinations resulted their fitness levels improved out of the sick/lethal range (fitness > -5), as observed both in our high throughput system (**Fig. 5F**) and by patch assay (**Fig. S8A**). We summarize all the observed interactions in **Fig. 5G**, representing the complicated higher-order epistasis network.

We also analyzed the epistasis networks formed by substitutions inthe *P. persalinus (Ciliate)* TL haplotype and identified two higher-order epistasis networks. The complexity of these networks arises from the intricate layers of substitution interactions. Notably, the complex interactions among these substitutions were not immediately apparent when examining the entire haplotype, as the expected and observed fitnesses of the complete haplotype were similar (**Fig. 6 and Fig. S9-S10**). First, the haplotype contains nine substitutions, each showing no or only slight growth defects as single substitutions. The observed fitness of the haplotype was very similar to the expected fitness calculated from the additive of all individual substitutions’ fitnesses, resulting in a primary deviation score close to zero. Second, when dissecting potential interactions in substitution combinations, we noticed distinct behaviors in two substitutions, the predicted GOF A1076T and LOF V1089S. We anticipated a suppression interaction between them and predicted that they would be drivers of epistatic interactions within intermediate combinations.

**Figure 6.**
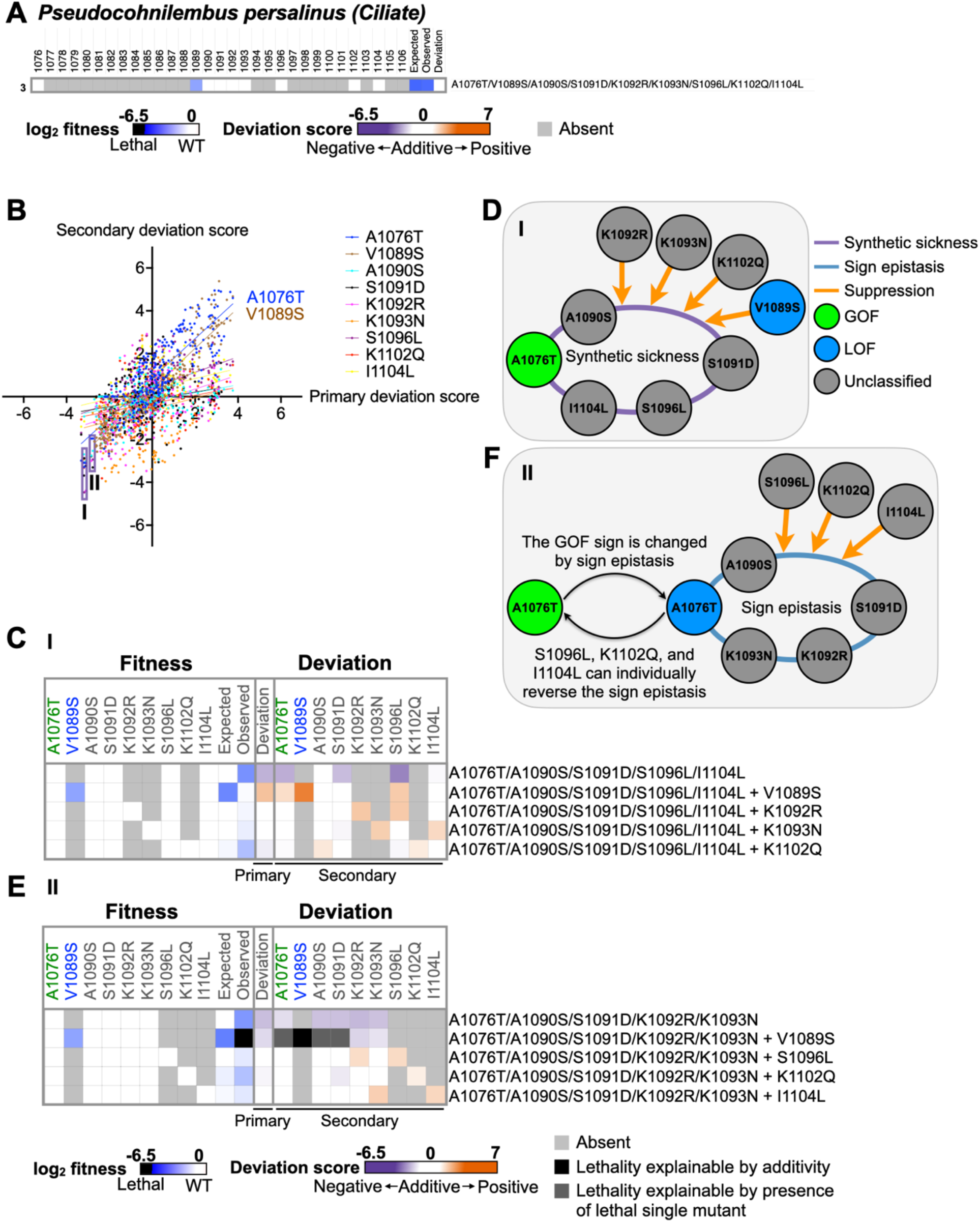
Intricate higher-order epistasis observed in substitutions of *P. persalinus (Ciliate)* TL haplotype. **A**. The heatmap displays the fitness of nine single substitutions in *P. persalinus (Ciliate)* TL in the yeast Pol II background, along with the epistasis between them represented by the primary deviation score. **B**. Similar to Fig. 5A, we checked correlations between secondary deviation scores (*y*-axis) to the primary deviation score (*x*-axis) to identify substitutions with substantial impact on primary deviation scores. Simple linear regression was applied to each comparison. Substitutions with R^2^ > 0.5 are annotated in the plot. **C**. The fitness and deviation scores of substitution combinations related to group I are shown in the heatmap. Names of substitutions are colored based on their phenotypic profiles as single mutants. GOF: green. LOF: blue. No obvious phenotype (unclassified): grey. Each line shows the fitness and deviation scores of substitutions in a certain combination. Left, the fitness of individual substitutions, and the expected and observed fitness. Right, the primary deviations calculated by comparing observed and expected fitness and the secondary deviation scores of each constituent substitution. A1076T/A1090S/S1091D/S1096L/I1104L is in the first line. Its observed fitness is smaller than expected and when compared, resulting in a negative primary deviation score, representing a negative interaction. The secondary deviation scores of each constituent substitutions are all negative, indicating each of them showing negative interactions when adding to corresponding compounds. The following four lines represent the four combinations where V1089S, K1092R, K1093N, and K1102Q are incorporated into A1076T/A1090S/S1091D/S1096L/I1104L respectively. All observed fitness of combinations is healthier than A1076T/A1090S/S1091D/S1096L/I1104L, and the secondary deviation scores of V1089S, K1092R, K1093N and K1102Q are all positive, implying positive effect (suppression) on each combination respectively. **D**. Scheme illustrating the substitution interaction network observed in C. **E**. Similar to C, the fitness and deviation scores of combinations related with group II are shown in the heatmap. The first row shows the fitness and deviation scores detected within the combination A1076T/A1090S/S1091D/K1092R/K1093N. The following rows displays the corresponding fitness and deviation scores when the other four substitutions are incorporated. Notably, the effect of V1089S on A1076T/A1090S/S1091D/K1092R/K1093N cannot be determined because the observed fitness of the combination (V1089S + A1076T/A1090S/S1091D/K1092R/K1093N) is in the ultra-sick/lethal range, and its expected fitness calculated from the log additive model is also in the lethal range due to additivity. In this case, the secondary deviation of V1089S is represented by a black block in the heatmap. Moreover, the effect of A1076T on V1089S/A1090S/S1091D/K1092R/K1093N cannot be determined because the observed fitness falls within the lethal range. The expected fitness of (A1076T + V1089S/A1090S/S1091D/K1092R/K1093N) is also in the lethal range due to the presence of the lethal compound V1089S/A1090S/S1091D/K1092R/K1093N. The expected lethality of the combination is because it contains a lethal component. The secondary deviation score of A1076T cannot be determined either and is indicated by a dark gray block in the heatmap. Similarly, the scores of A1090S and S1091D could not be determined for the same reason with A1076T and are also marked with dark grey blocks. **F**. Scheme representing the substitution interaction networks observed in E.

Consistently, we observed suppression when A1076T and V1089S were combined (**Fig. S9**), and correlations between the secondary and primary deviation scores suggest that these two substitutions strongly affect the epistasis in the haplotypes (**Fig. 6B**). Third, we identified two intermediate combinations, each with five substitutions, that exhibited the lowest primary and secondary deviation scores (the labeled **I** and **II in Fig. 6B**). These low scores imply negative interactions such as synthetic sickness or lethality.

The combination I, A1076T/A1090S/S1091D/S1096L/I1104L, showed a negative primary deviation score (observed fitness < expected fitness) (**Fig. 6C**), illustrating synthetic sickness among the five substitutions. The secondary deviation scores of five constituent substitutions were all negative (**Fig. 6C**), implying that each addition worsened the fitness. Conversely, the remaining four substitutions, V1089S, K1092R, K1093N, and K1102Q, all showed positive secondary deviation scores when incorporated into the combination. This suggests each one could individually suppress the synthetic sick combination (**Fig. 6C**). We validated the suppression by V1089S to the combination by spot assay and doubling time (**Fig. S8B-C**). The consistency indicates the interactions we observed in the high throughput experiment are reliable. The observed interaction network is shown in **Fig. 6D**.

In contrast, combination II (A1076T/A1090S/S1091D/K1092R/K1093N) exhibited sign epistasis, where the GOF sign of A1076T appeared to change to LOF in the presence of the other four substitutions simultaneously. In detail, the combination II has a negative primary deviation score, representing negative interaction among five substitutions (**Fig. 6E**). Surprisingly, the negative interaction appeared only when the four substitutions, A1090S, S1091D, K1092R, and K1093N were present simultaneously with A1076T (**Fig. S10A**). Notably, no obvious interactions occur among these four substitutions alone (**Fig. S9**), suggesting the interaction is between A1076T and the combination of the four substitutions (A1090S/S1091D/K1092R/K1093N). We further inferred a change in A1076T from GOF to LOF, particularly due to the observed lethality when LOF V1089S was incorporated into the combination. Notably, while A1076T generally suppresses V1089S in almost all synthetic sick or lethal backgrounds involving V1089S and the individual or combinations of A1090S, S1091D, K1092R, and K1093N, it exacerbates the growth defect when combined with V1089S/A1090S/S1091D/K1092R/K1093N. This outcome aligns with the expected lethality when the same class substitutions are combined (LOF V1089S and the LOF A1076T) (**Fig. S10B-C**). These observations suggest that the combination A1090S/S1091D/K1092R/K1093N converts A1076T from GOF to LOF. Further supporting this, when S1096L, K1102Q, and I1104L are individually included in the combination A1076T/A1090S/S1091D/K1092R/K1093N, each one suppresses the combination, as evidenced by their positive secondary deviation scores (**Fig. 6E**). The weak suppression between I1104L and the combination was also confirmed by comparison in doubling time (**Fig. S8C**). The suppression interactions imply that each one of them can individually revert the sign epistasis within A1076T/A1090S/S1091D/K1092R/K1093N. As a result, A1076T is back to GOF, and LOF V1089S becomes suppressible by GOF A1076T (**Fig. 6F**). In summary, these intricate interaction networks observed in the *P. persalinus (Ciliate)* TL haplotype (**Fig. 6E-F**) emphasize the complexity of higher-order epistasis, revealing layers of interactions beneath the surface.

### Distinct categories of residue epistasis patterns

To comprehensively compare the magnitude and consistency of epistatic effects of TL substitutions across different genetic backgrounds, we determined the distributions of secondary deviation scores for each substitution. Substitutions with strong epistatic effect (**Fig. 4 and S6**) and primary/secondary score correlations (**Fig. 5A**, **Fig. 6B, and Fig. S7**) exhibited wide distributions of secondary deviation scores. Interestingly, while most of these impactful substitutions have classifiable phenotypes as single substitutions (GOF or LOF) (**Fig. S11A-F**), others without obvious phenotypes, such as S1091E (**Fig. S11A**), K1093N (**Fig. S11E-F**), A1076G and N1082K (**Fig. S11G**), still showed wide ranges of epistatic impacts. Notably, some substitutions, like A1076T, were present in more than one TL haplotype background (**Fig. S11C-F**). To evaluate the overall epistatic effects of these substitutions in all tested backgrounds, we displayed the density plots of their secondary deviation scores across all non-repetitive haplotypes (**Fig. S12**) and calculated the maximum likelihood estimate (σ^2^) to quantify the distribution of secondary deviation scores for each substitution (PARK *et al*. 2022) (**Fig. 7A**). We found that 62.5% of substitutions exhibited mild epistatic effects (σ^2^ < 3) and, 25% showed medium effect (3 ≤ σ2 ≤ 5). A small portion, 12.5% of substitutions, had strong epistatic effects (σ^2^ > 5). The epistatic effects of substitutions did not correlate with their fitness as single substitutions (**Fig. S13A**). Moreover, if the substitution has no or minor epistatic effect in backgrounds to which it is introduced, its secondary deviation scores are expected to follow normal distribution (PARK *et al*. 2022). However, approximately 80% of TL substitutions did not have normally distributed secondary deviation scores (**Fig. S13B**), illustrating that most TL substitutions had epistatic effects when introduced into some if not all genetic backgrounds. To investigate whether the epistatic effects of substitutions consistently remained positive (or negative) across various genetic backgrounds, we calculated the median of secondary deviation scores for each substitution and plotted them against the corresponding σ^2^ values. Substitutions with strong epistatic effects (σ^2^) exhibited higher positive (or negative) median values (**Fig. 7B**), implying that robust epistatic effects were consistently positive or negative across various genetic backgrounds. We further compared the epistatic effects of substitutions in three categories (**Fig. 7C**). Substitutions with mild epistatic effects had little impact on the fitness of haplotypes when introduced. Substitutions with medium epistatic effects, like A1076T and K1093N, could either enhance or reduce haplotype fitness. In contrast, substitutions with strong epistatic effects, such as S1096E and R1100C, were epistatic to backgrounds in which they were observed, consistent with their requirement for tolerance of specific substitutions in those backgrounds.

**Figure 7.**
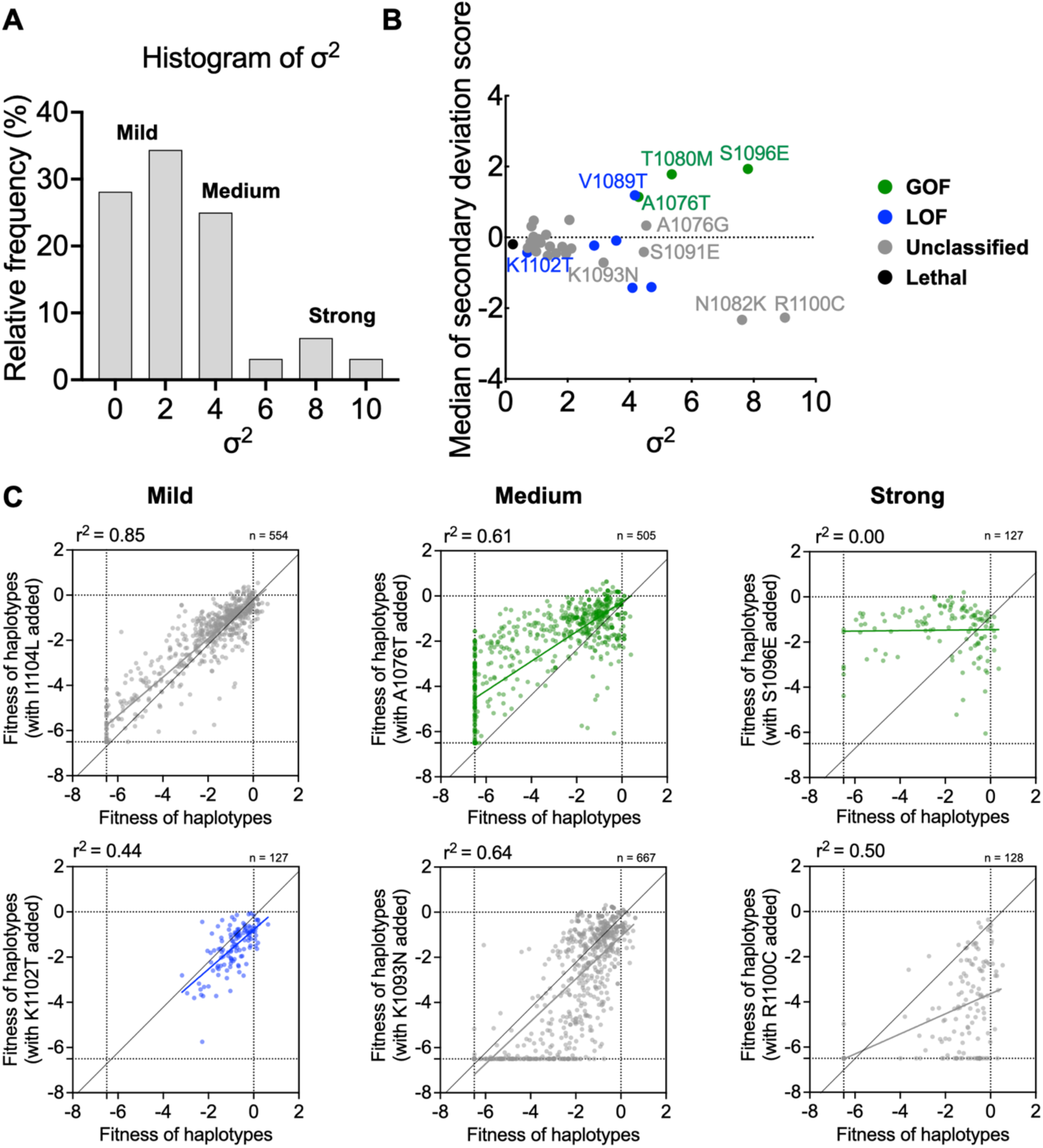
Different classes of epistatic effects. **A**. Histogram of mutants’ epistatic effects, represented by their respective maximum likelihood estimate (σ^2^) of secondary deviation scores. Higher epistatic effect indicates greater impact of a certain substitution. **B**. Medians of secondary deviation scores of substitutions were plotted against their corresponding σ^2^. Substitutions are colored based on their phenotypes. **C**. Comparing epistatic effects of mutants in each category. Each scatter plot shows the measured fitness of haplotypes without (*x*-axis) versus with (*y*-axis) a substitution incorporated. The colors of the plots represent the mutants’ phenotypes. The colored line marks the simple linear regression of the spots, representing the observed epistatic effect of the substitution. R^2^ values of the regressions are labeled in the plots. The black line indicates the additive (non-epistatic) expectation.

## DISCUSSION

RNA polymerase TL function and evolution are impacted by extensive residue interactions within and around it (KAPLAN *et al*. 2012; DUAN *et al*. 2023). The TL shows remarkable conservation, consistent with its essential role in executing transcription mechanisms in all msRNAPs (CRAMER 2002; WERNER AND GROHMANN 2011; BELOGUROV AND ARTSIMOVITCH 2019; MAZUMDER *et al*. 2020; KULDELL AND KAPLAN 2024). Our observation of widespread TL incompatibility in the *S. cerevisiae* Pol II context suggested that epistasis external to the TL is the major source of epistasis affecting TL function (DUAN *et al*. 2023) (**Fig. S3**). However, here we observed TL-internal epistatic interactions and residue coupling, although TL-internal coupling did not significantly maintain TL function when the interaction between TL and its broader evolutionary context was disrupted. The analyses in our recent work (DUAN *et al*. 2023) and here indicate that interactions formed by TL residues are integrated into the broader residue epistasis network within the whole enzyme (**Fig. 2**). These results align with other studies showing that the same mutation led to different phenotypes when introduced into homologous proteins (KONDRASHOV *et al*. 2002; LUNZER *et al*. 2010; NATARAJAN *et al*. 2013; VIKTOROVSKAYA *et al*. 2013; DOUD *et al*. 2015; STARR AND THORNTON 2016; HADDOX *et al*. 2018; PARK *et al*. 2022), highlighting the divergent nature of epistasis networks even in highly conserved proteins like msRNAPs.

Our experiments here provide case studies for examples of complex interactions during evolutionary divergence. Epistasis networks of homologs diverge through accumulated substitutions. Understanding how substitutions alter existing epistasis and shape the effects of future substitutions is critical for elucidating mechanisms underlying protein functional divergence and evolution (STARR AND THORNTON 2016; JOHNSON *et al*. 2023; XIE *et al*. 2023). Using seven selected Pol II TL variants, we comprehensively determined the higher-order epistatic interactions created by these substitutions. For example, S1096E exhibited strong suppression on all synthetic sick/lethal combinations in the *E. invadens IP1* TL haplotype (**Fig. 5 and S8**), suggesting a potential evolutionary pathway for TL where the 1096E substitution occurs prior to additional substitutions that lead to synthetic sickness or lethality. A caveat to this interpretation is that additional substitutions outside the TL might reshape epistasis across the entire TL. Additionally, A1076T, a substitution that consistently showed positive epistatic effects when introduced into different genetic backgrounds (**Fig. 7C**), was also subject to sign epistasis in certain backgrounds. Here, we would propose that additional substitutions preventing this sign epistasis might precede the appearance of combinations that would otherwise incur a fitness defect with A1076T (**Fig. 6E-F**). The sign epistasis involving A1076T exemplifies the intricate layers of substitution interactions, where emerging substitutions modify pre-existing interactions, ultimately influencing the fate of future mutations and the trajectory of protein evolution.

We further inquired whether individual substitutions could alter pre-existing epistasis networks in the protein background they were introduced into, and to what extent. To address this, we analyzed the strength and consistency of the epistatic effects of substitutions across diverse genetic backgrounds. TL substitutions can be categorized into three groups based on their epistatic effects. Approximately 37.5% of TL substitutions consistently demonstrated medium to strong, stable epistatic effects, either positive or negative (**Fig. 7A-B**), reflecting their important role in reshaping the interactions among fixed historical substitutions and influencing the phenotypes of upcoming mutations. Substitutions that consistently exhibit positive epistatic effects, such as A1076T and S1096E (**Fig. 7C**), are referred to as “permissive substitutions” (STARR AND THORNTON 2016). They have the potential to make certain mutations accessible that would otherwise remain inaccessible. An obvious path for evolutionary divergence in msRNAPs would be the incorporation of substitutions with mild effects on fitness but with biochemical properties of increased catalytic activity. These GOF alleles would have properties of suppression of LOF alleles that might otherwise be selected against. Conversely, substitutions with a consistent negative effect, like K1093N and R1100C, may act as “restrictive substitutions”, limiting the accessibility of some mutations (STARR AND THORNTON 2016). A limitation of our analyses is that it tested a limited number of backgrounds. In the future, our platform has the capability to determine if the case studies presented here are rarer or the norm in the TL. In summary, our analyses provide a framework for understanding complicated epistatic interactions of substitutions and examples of how fitness landscapes of mutants are changed due to epistasis.

## METHODS

### Experimental data

Experimental data was collected as described in Duan et al (DUAN *et al*. 2023). Briefly, we synthesized TL mutant libraries including 632 natural TL variants from 182 bacterial, 116 archaeal, and 333 eukaryotic RNA polymerases (Pol I TLs n = 144, Pol II TLs n = 94, Pol III TLs n = 95) to detect their compatibility in *S. cerevisiae* background. Additionally, to identify internal interactions within TL haplotypes, we selected seven Pol II TL natural variants from eukaryotic Pol II (names of the selected species are as described in **Fig. 3B-H**), and synthesized 1987 TL haplotypes encompassing every possible substitution combination among the seven selected Pol II TL haplotypes. Furthermore, we designed and synthesized 724 Pol II TL haplotypes specifically for coupling analysis. Lastly, 620 TL single mutants serving as control to determine residue interactions. Approximate 15% wild-type *S. cerevisiae* Pol II TL allele of total variants was incorporated into each TL library for accurate quantification. All TL alleles were amplified and transformed into yeast strain CKY283 along with modified *RPO21/RPB1*-encoding plasmid to allow construction of complete *rpb1* TL mutants within yeast through gap repair. After transformation, Leu+ colonies were collected, and re-plated into subsequent selection plates (SC-Leu+20mg/L Mg, SC-Leu+15mM Mn, YPRaff, YPRaffGal, SC-Lys, and SC-Leu+3% Form).

Growth defects and phenotypes of mutants were assessed through amplicon sequencing with Illumina Next-seq (150nt reads). We carefully constructed amplicon sequencing libraries. We performed emulsion PCR on the TL region to amplifly the mutants in the libraries while minimize template switching. Additionally, PCR was done with optimized cycles to preserve original allele frequencies. Subsequently, we employed dual indices for multiplexing samples.

### Data cleaning and normalization

Each TL variant library was constructed and subject to screening in three biological replicates. After screening, TL regions of mutants in the variant libraries were amplified and sequenced. The read counts of mutants were estimated by a codon-based alignment algorithm (SING-HOI SZE 2018), and the median of these counts from the three replicates was used for calculating growth defects in subsequent analysis. Growth defects of mutants were represented with a fitness metric. Fitness is determined by comparing the shifts in allele frequency of a mutant under selective conditions compared to control conditions, with these shifts further compared against changes observed in the WT. The log of the comparison is defined as a fitness. The formula used for this calculation is detailed below.

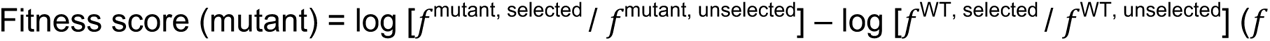

represents allele frequency of a certain variant in this formula).

To enable direct comparisons between libraries, we applied min-max normalization to standardize median growth fitness across all variants. Residue with fitness equal or smaller to -6.5 was categorized within the severe sick and lethality range, and their fitness are normalized to the lethal threshold of -6.5, ensuring consistency in the severity classification.

### Determination of functional interactions

Functional interactions were represented by genetic interactions, calculated using the log additive model(**Fig. S1B**). This model assumes individual mutations act independent, and their effects are multiplied in the double/multiple mutants. In our case, the fitness scores are expressed in logarithmic form, so these effects become additive.

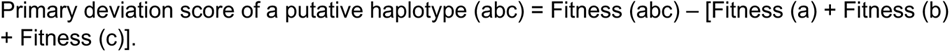

We further selected seven Pol II TL alleles and constructed all intermediate substitution combinations. For example, the putative TL haplotype has three different residues compared with wild-type *S. cerevisiae* Pol II TL allele a, b and c, we constructed all combinations of substitutions, which are a, b, c, ab, ac, bc, and abc. And we can calculate the secondary deviation scores for each substitution a, b and c. The secondary deviation score of a are calculated from all haplotypes containing a: Secondary deviation score of a_1_ = Fitness (ab) – Fitness (b). Secondary deviation score of a_2_ = Fitness (ac) – Fitness (c). Secondary deviation score of a_3_= Fitness (abc) – Fitness (bc).

Primary and secondary deviation scores cannot be determined if the expected and observed fitness are both in the sick/lethal range (both fitness < -6.5). Haplotypes with expected fitness < -6.5 can be separated into two different situations. First, when all constituent substitutions are viable, but the sum of the fitness is lower than -6.5. This would be consistent with lethality due to additivity and indicated with a dark gray block in the deviation score heatmaps. Second, when some of the constituent substitutions or combinations are in lethal range, this lethality is reasoned to derived from the haplotype containing at least one individually lethal substitution and indicated with a black block in the deviation heatmap.

Functional interactions are determined from deviation scores. Positive interaction (suppression): deviation score > 1. Additive (No interaction): -1 ≤ deviation score ≤ 1. Negative interaction (synthetic sickness/lethality): deviation score < -1.

Maximum likelihood estimate (σ^2^) of a certain substitution is calculated to represent its overall epistatic effect (PARK *et al*. 2022). For a substitution with n non-repetitive secondary deviation scores, its maximum likelihood estimate: 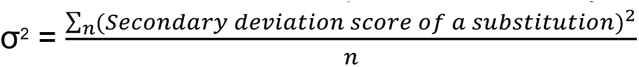

### Coupling analysis

Coupling analysis was done as described in Russ et al (RUSS *et al*. 2005) and Socolich et al (SOCOLICH *et al*. 2005) using a multiple sequence alignment (MSA) of Pol II TL sequences. Briefly, three independent TL haplotypes libraries were constructed to study intra-TL co-evolutionary residues. (a). Natural variants library, which consists of 362 selected natural eukaryotic Pol I, II, and III TL variants. To detect potential coupled residues, these TL variants were aligned using multiple sequence alignment, followed by coupling analysis. (**Fig. S3A**). (b). Random scrambling library, where amino acids in the same position of the MSA from (a) were randomly swapped. This swapping does not disturb sequence conservation, because the composition of different amino acids in the position remains unchanged but disrupts residue coupling among different residues. Coupling analysis was done for the random scrambled MSA to confirm disrupted residue coupling (**Fig. S3B**). (c). Monte Carlo scrambling library, where the amino acids in the same position were swapped and then assessed by the Monte Carlo algorithm. If the swapping perturbs residue coupling, we withdraw the swap; if not, we keep the swap. By doing this, the Monte Carlo scrambling library retains both sequence conservation and residue coupling information (**Fig. S3C**). The swapping in both cases breaks the epistasis between TL residues and the Pol II context. TL haplotypes were synthesized based on three types of MSA and then screened for mutant phenotypes through phenotypic system (QIU *et al*. 2016; QIU AND KAPLAN 2019; DUAN *et al*. 2023).

### Doubling time measurement

TL variants were tested in four biological replicates, each with three or four biological replicates. Overnight cultures of all variants were diluted to an OD of 0.005 in 200 μL YPD and grown at 30°C, with OD measurements taken every 30 minutes for 48 hours. The experiment was done using a Tecan Spark Microplate Reader. R package Growthcurver (version 0.3.1) was used in analysis for obtaining doubling time (SPROUFFSKE AND WAGNER 2016).

## DATA AVAILABILITY

Raw sequencing data has been deposited on the NCBI SRA (Sequence Read Archive) database under BioProject PRJNA948661. Processed mutant count, fitness and processing codes are available through GitHub (https://github.com/Kaplan-Lab-Pitt/TLs_Screening.git). Strains and plasmids will be provided upon request.

## AUTHOR CONTRIBUTION

BD performed sequencing library construction, data analysis, and made figures, drafted and revised the manuscript. CQ designed the mutant libraries, performed screening experiments and statistical coupling analysis, and contributed to revise the manuscript. SZ counted the mutants from raw sequencing data. SWL contributed to experimental design and manuscript revision. CDK conceived the project, guided analyses and interpretation of data, provided funding, revised the manuscript.

## ACKNOWLEDGMENTS

We thank Dr. Anne-Ruxandra Carvunis (U. Pittsburgh) for discussions and advice. We thank Zhizhen Wang and Muyao Lin from the Pitt Statistical Consulting Center for their advice on checking the reproducibility of our data.

---

**Figure S1.**
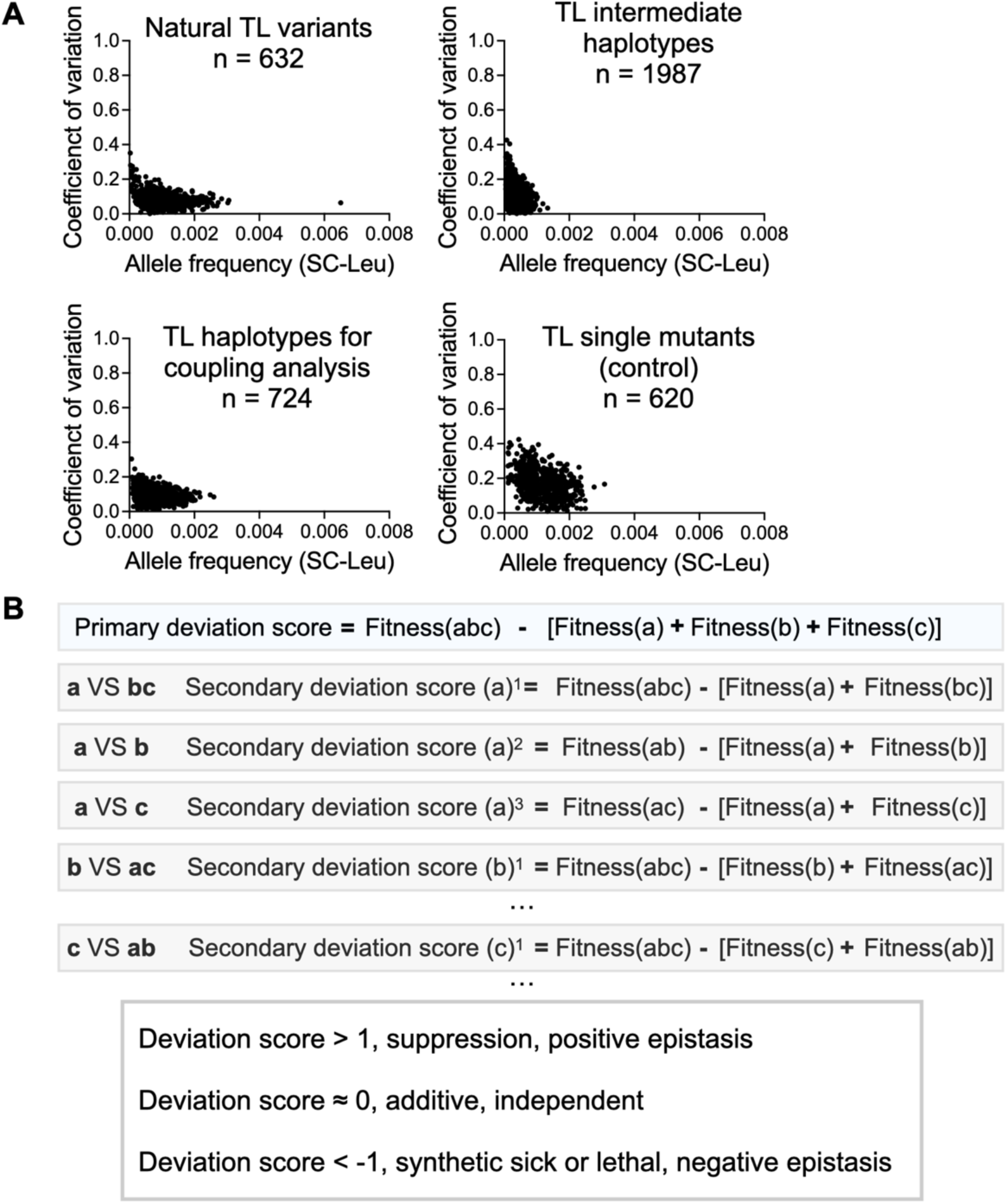
Detection of higher-order interactions by primary and secondary deviation scores. **A**. The low coefficients of variation for all libraries involved in this study indicate high reproducibility across the datasets. The coefficient of variation is plotted against the allele frequency in the control condition, SC-Leu, for each of the four libraries. **B**. For a pseudo haplotype “abc”. The primary deviation score is calculated by comparing the observed fitness of the haplotype to the log additive of constituent substitutions’ fitness. The secondary deviation score of “a” in “abc” represents the epistatic effect of “a” on “bc” and is calculated by comparing the observed fitness score of “abc” to the additive of fitness scores of “a” and “bc”. The secondary deviation score of “a” in “ab” represents the epistatic effect of “a”on “b”. It is calculated by comparing the observed fitness score of “ab” to the additive of fitness scores of “a” and “b”. If the deviation score ≈ 0, it indicates the constituent substitutions are independent and there is no interaction among them. If the deviation score > 1, it represents positive epistasis (suppression) among the constituent substitutions. If the deviation score < -1, it represents negative epistasis (synthetic sickness or lethality) among the constituent substitutions.

**Figure S2.**
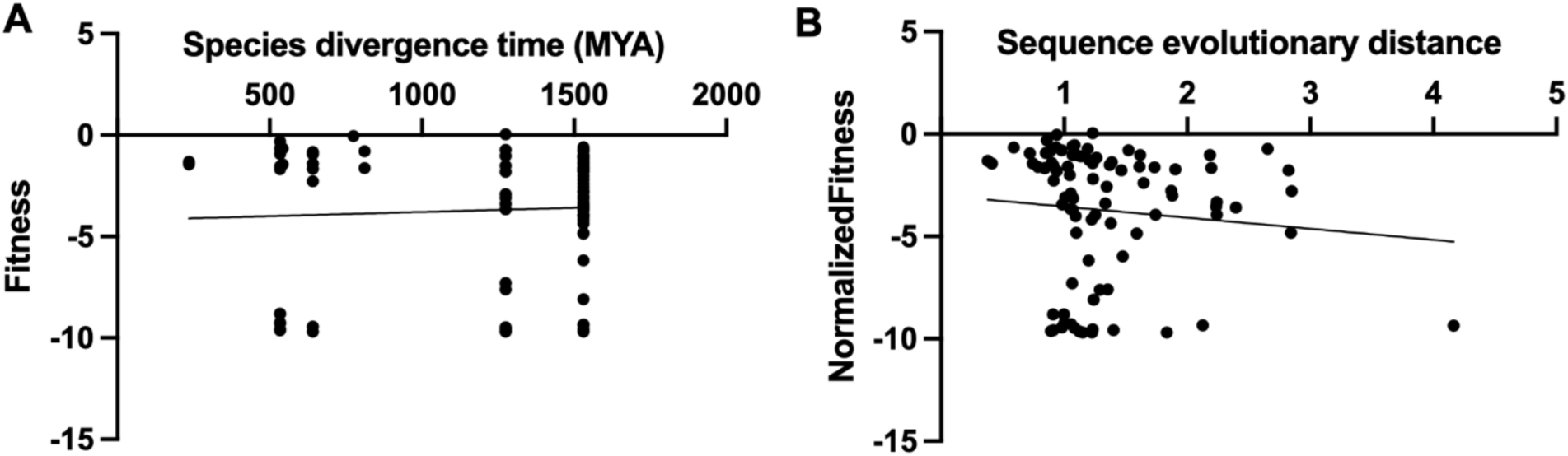
Evolutionary distances do not correlate with the growth fitness of TL haplotypes. **A**. The divergence time between the species from which the TL haplotypes originate and the yeast *Saccharomyces cerevisiae*, which contains the WT TL, does not correlate with the fitness of the Pol II TL haplotypes. The median divergence times showing in axis are represented with millions of years ago (MYA) and calculated by TimeTree5 (KUMAR *et al*. 2022). The line represents the simple linear regression between the divergence time and the fitness: *y* =0.0004123\**x*-4.201 (slope is not significantly different from zero). **B**. The evolutionary distance between the genes that the Pol II TL haplotypes are from and the yeast *RPB1* gene, which contains the WT TL, does not correlate with growth fitness. The sequence evolutionary distances are calculated by MEGA11 with JTT model (TAMURA *et al*. 2021). The line represents the simple linear regression between the evolutionary distance and the fitness: *y* = -0.5401 \**x* – 3.005 (slope is not significantly different from zero).

**Figure S3.**
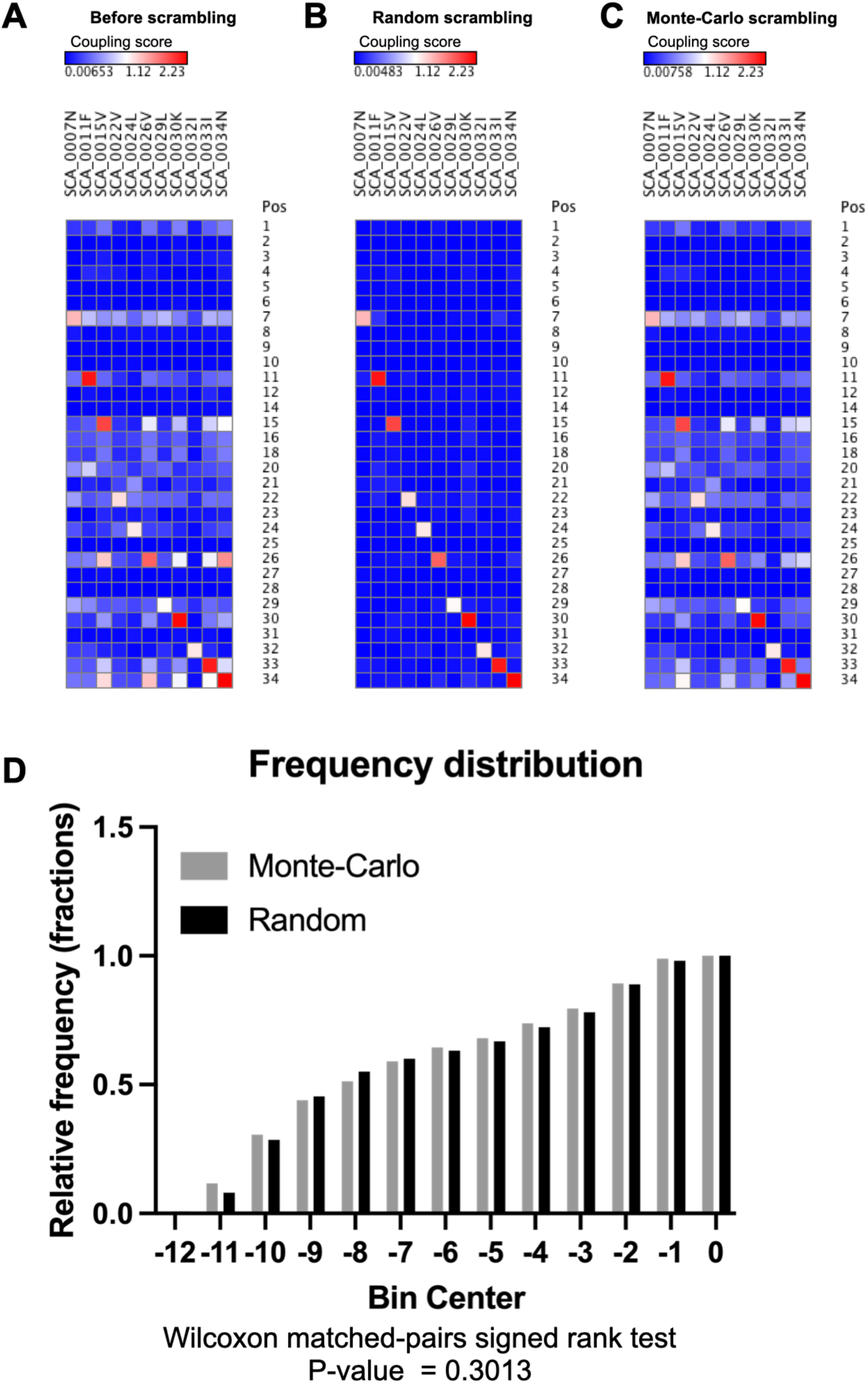
Design of mutant libraries to assess TL-internal residue coupling. **A**. Residue coupling heatmaps of natural TL variants. Statistical coupling analysis of TL multiple sequence alignment including 362 selected natural eukaryotic Pol I, II, and III TL alleles. **B-C**. Heatmaps of coupled residues in Random scrambling and Monte Carlo scrambled haplotypes. The random scrambling disrupted coupled residues in selected TL variants (**B**) while the Monte Carlo scrambling haplotypes reserved coupled residues (**C**). **D**. Cumulative fitness frequency distribution of the Random scrambling and Monte Carlo scrambling haplotypes. The Wilcoxon matched-pairs signed rank test was used to assess the significance of the fitness distribution difference between Monte Carlo scrambling and Random scrambling haplotypes. The P-value indicates that they are slightly but significantly different.

**Figure S4.**
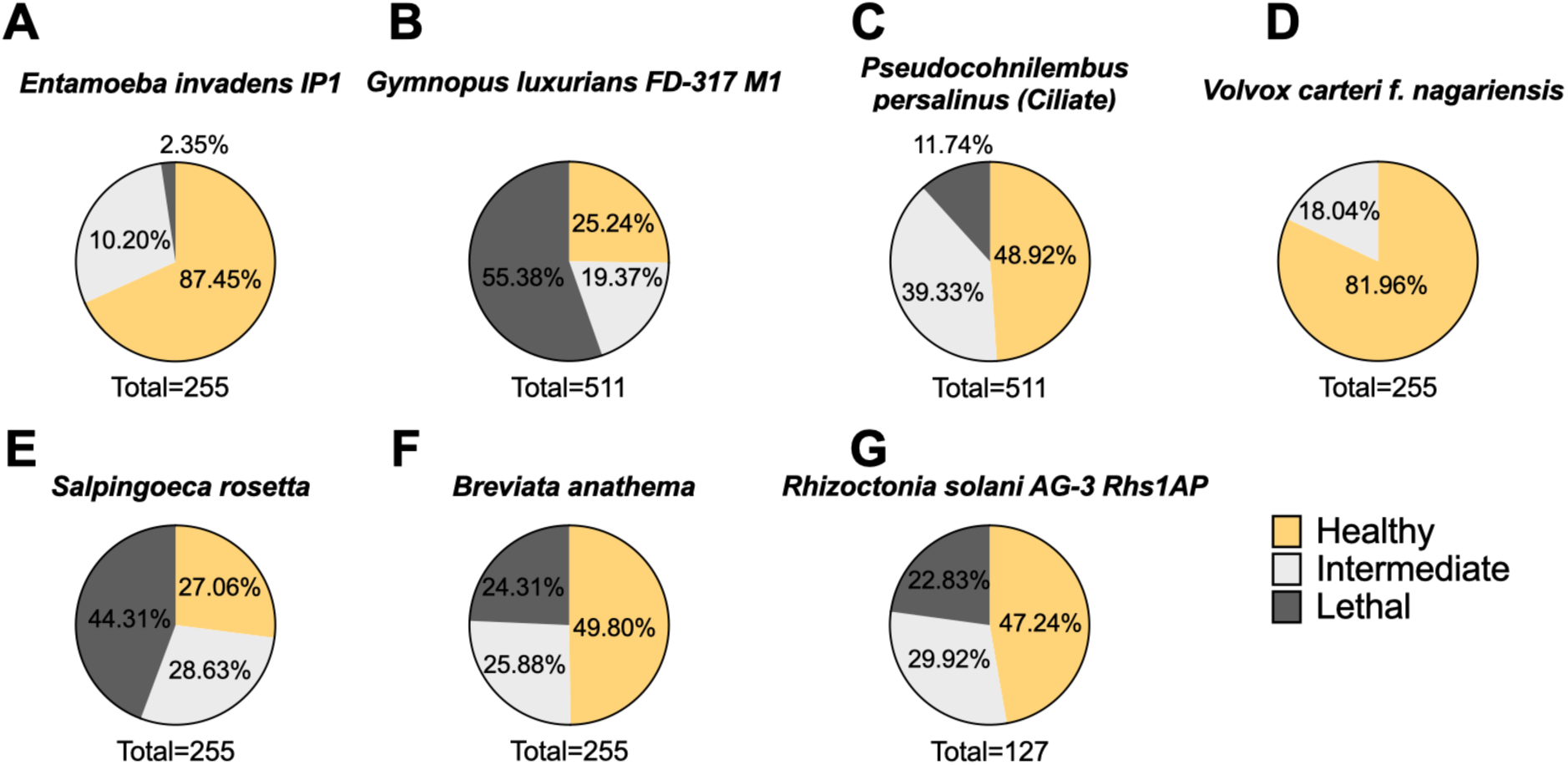
The percentage of fitness categories of all intermediate combinations within each haplotype. **(A-G)**. Healthy: Fitness > -2. Intermediate: -6.5 < Fitness ≤ -2. Lethal: Fitness ≤ -6.5.

**Figure S5.**
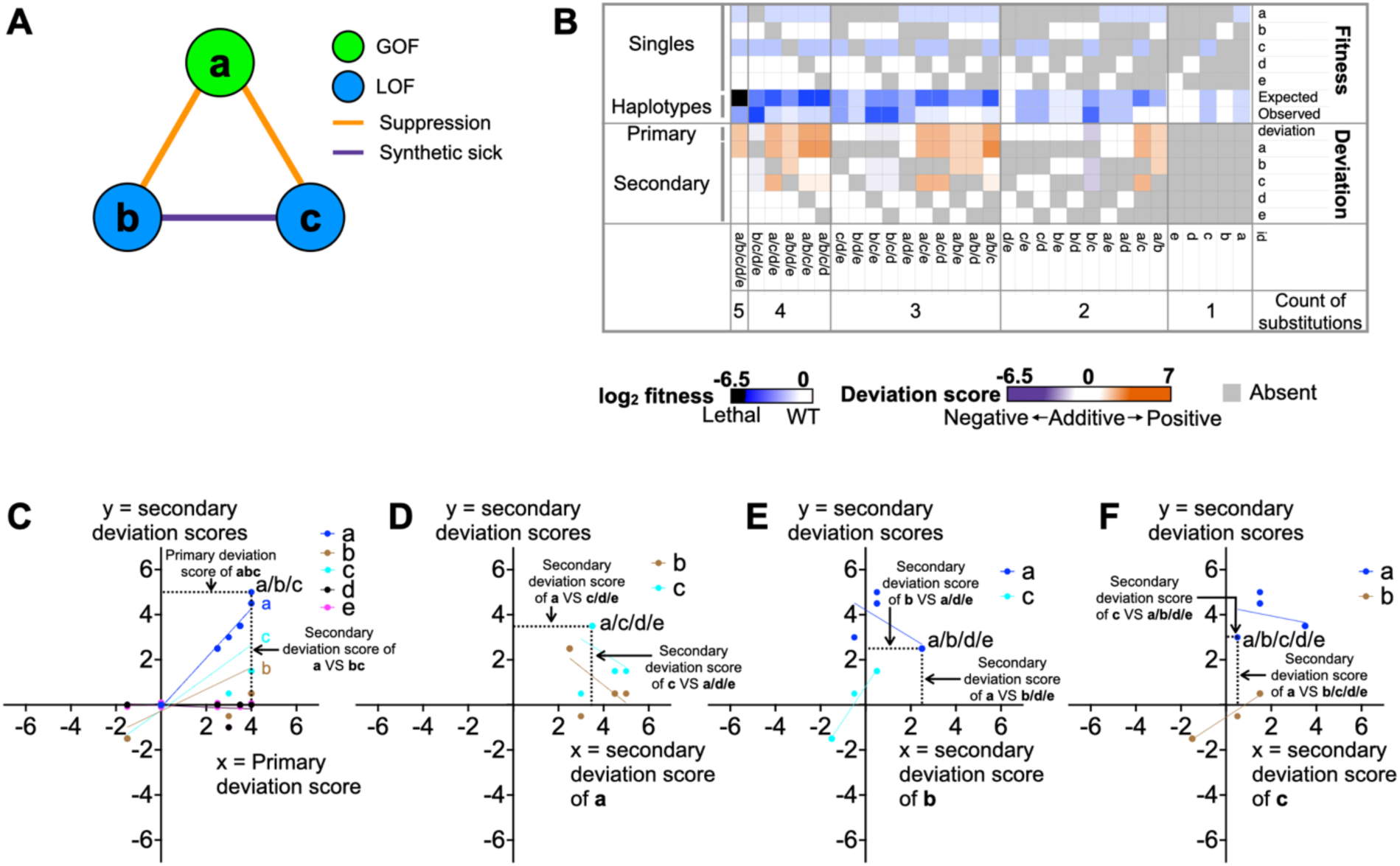
Example primary and secondary deviation scores for a simplified, simulated TL haplotype “abcde”. **A**. Simulated interactions among substitutions “a”, “b”, and “c”. We set the single mutant “a” as a GOF mutant, “b” and “c” as LOF mutants, while “e” and “f” have no effects. We also set suppression between GOF and LOF combinations (“ab” and “ac”), and synthetic sickness between LOF and LOF combinations (“bc”). **B**. The epistasis interaction landscape of simplified TL haplotype “abcde”, illustrating the observed and expected fitness of all single substitutions and all intermediate haplotypes, as well as primary and secondary deviation scores in the simulation. **C**. Correlations between secondary deviation scores of all five single substitutions (*y*-axis) to the corresponding primary deviation scores (*x*-axis). To measure the correlation, linear regression was applied for secondary deviation scores of “a”, “b”, “c”, “d”, “e” to the corresponding primary deviations scores, respectively. Single substitutions with strong correlations (R^2^ > 0.5) were labeled in the plot. Three substitutions with phenotypes show strong correlations in the simulation, “a”, “b”, and “c”. **D-F**. The correlation between secondary deviation scores of the four substitutions (*y*-axis) vs “a” (**D**), “b” (**E**), “c” (**F**) on *x*-axis respectively. To read the plot, with **D** as an example, each cyan spot represents an intermediate combination containing both “a” and “c”. The labeled spot represents a specific combination “a/c/d/e”. Its coordinate value in *x*-axis represents the secondary deviation score of “a” to “c/d/e”. Its coordinate value in *y*-axis represents the secondary deviation score of “c” to “a/d/e”. We did a linear regression for all the cyan spots to represent the correlation between the secondary deviation scores of “c” to “a”. This correlation helps illustrate the potential interaction between “c” and “a”. The secondary deviation scores of “c” are all positive when “a” is present, meaning “c” constantly shows positive secondary score when “a” is present, indicating potential suppression of “c” to “a”.

**Figure S6.**
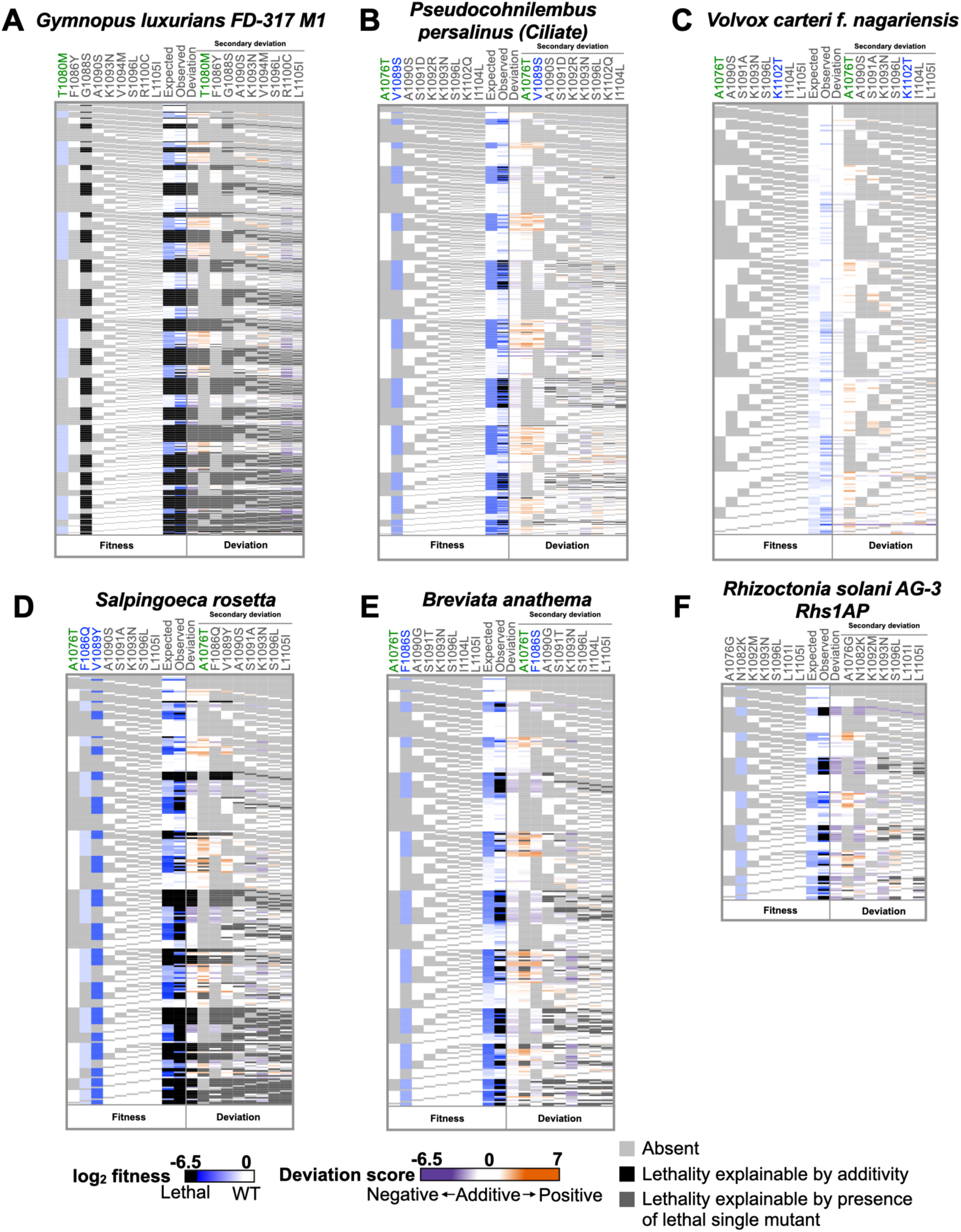
The epistasis landscapes of selected haplotypes. The fitness and epistasis of all unique intermediate haplotypes from combinations of substitutions within each haplotype (**A-F**). The colors of mutants’ names represent mutants’ phenotypes. GOF is in green, LOF is in green, non-classified mutants is in grey.

**Figure S7.**
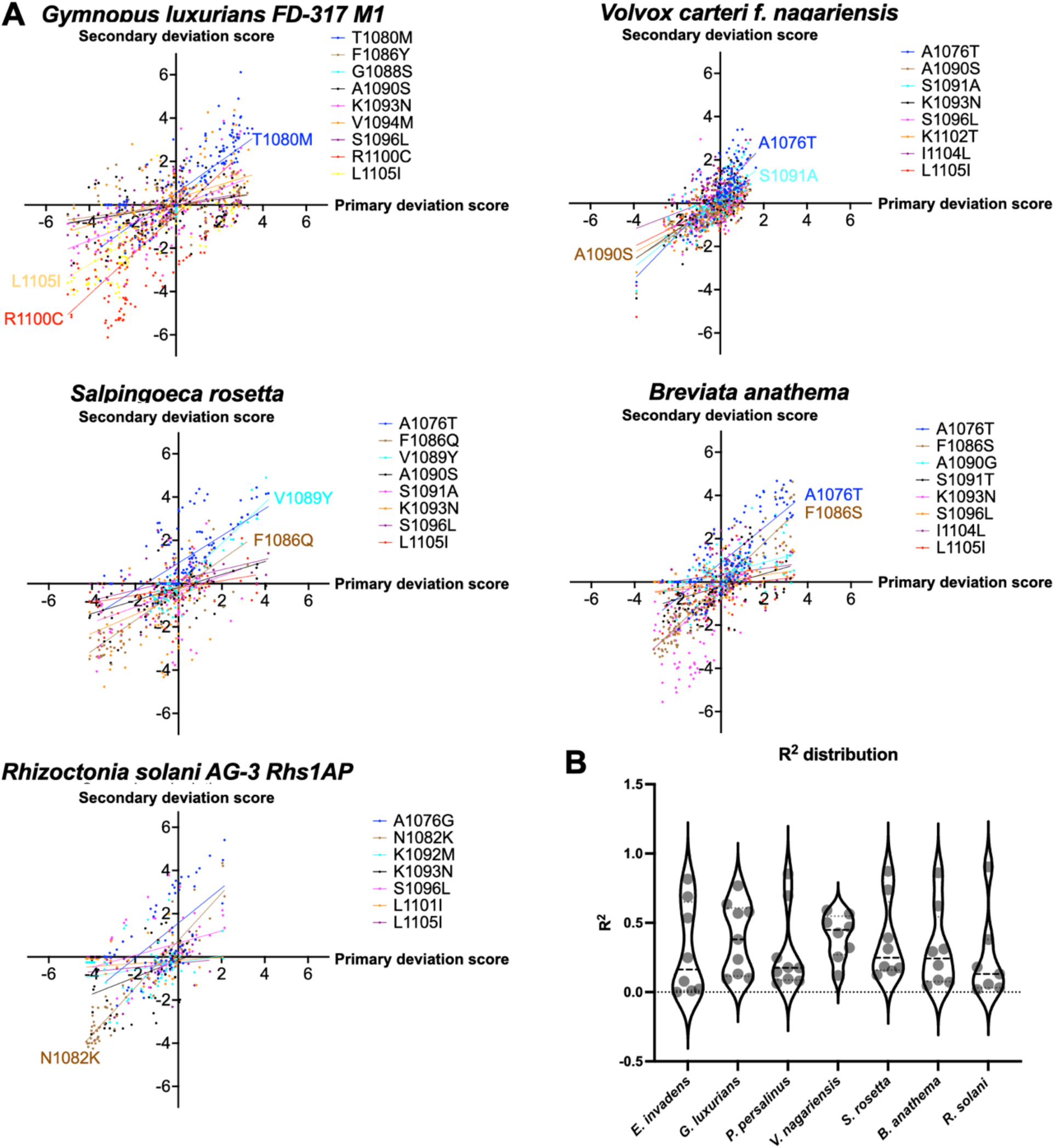
The epistasis within haplotypes is driven by several specific substitutions. **A**. The correlations between secondary deviation scores of all substitutions (*y*-axis) and the primary deviation score (*x*-axis) for each selected haplotype. Linear regression was applied to each comparison of secondary to primary deviation scores. Substitutions with R^2^ value exceeding 0.5 are annotated on the *x*-*y* plot, indicating their substantial impact on primary epistasis of the haplotypes. **B**. Distributions of R^2^ of all selected TL haplotypes.

**Figure S8.**
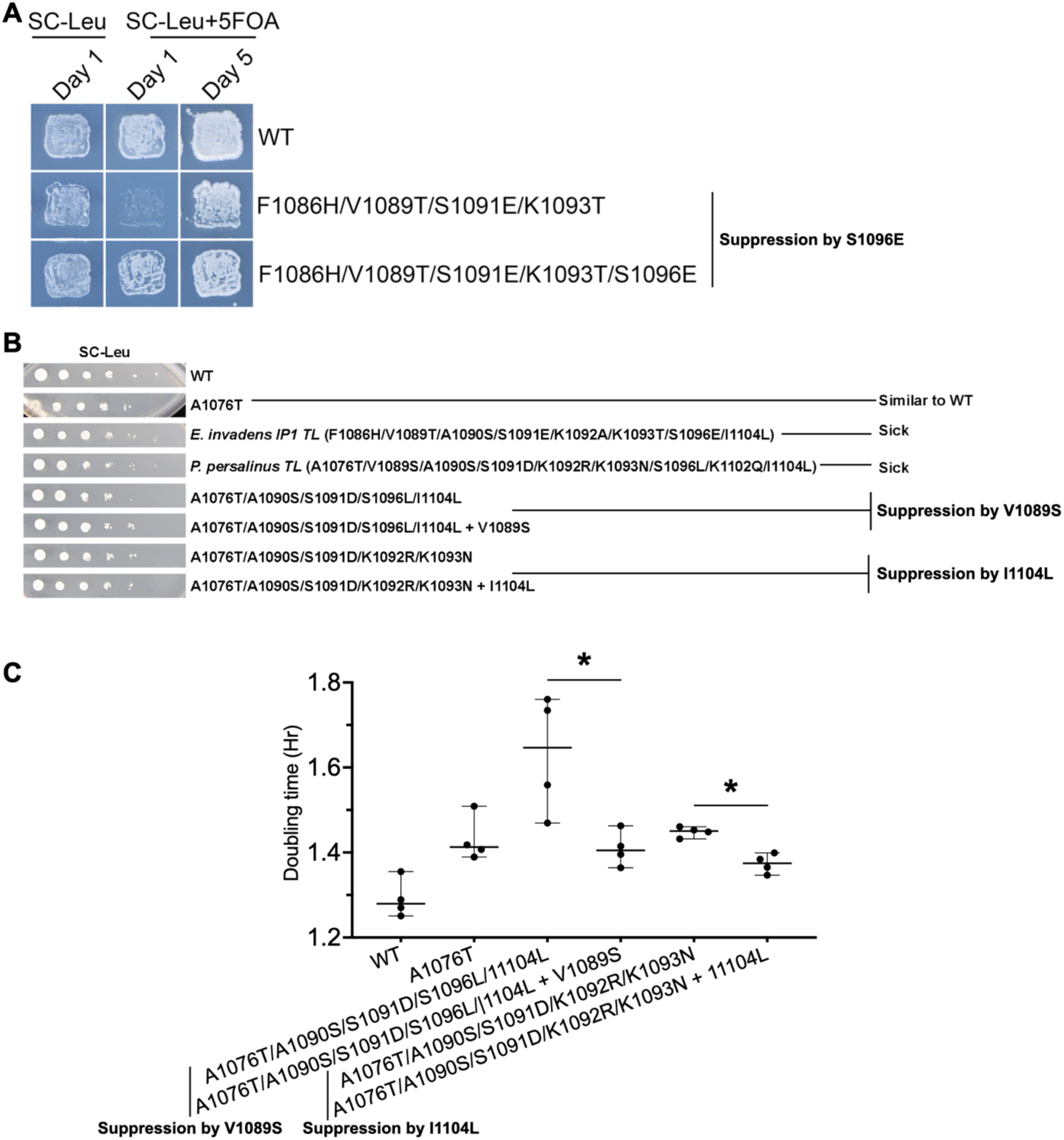
Validation of genetic interactions. Validating genetic interactions observed in high throughput libraries with growth assays. Haplotypes involved here were synthesized and transformed into WT yeast cells, and their sequencies were extracted and verified by sequencing. **A**. Confirmation of the suppression between S1096E and the synthetic sick/lethal group F1086H/V1089T/S1091E/K1093T by patch assay. Growth of patches on SC-Leu+5FOA indicates the growth defects of the haplotypes. F1086H/V1089T/S1091E/K1093T barely grew on SC-Leu+5FOA condition on Day 1, while F1086H/V1089T/S1091E/K1093T/S1096E showed near WT level growth, suggesting the suppression between S1096E and F1086H/V1089T/S1091E/K1093T. **B**. Verification of growth defects and genetic interactions in haplotypes and combinations by spot assay. The growth defects of *E. invadens* IP1 TL and *P. persalinus* TL were confirmed by their smaller spot sizes compared to the WT, consistent with our observations from the high throughput experiment. To validate the suppression interactions observed in the high throughput data, we selected two pairs of haplotypes. While suppression was detected in the spot assay, the interactions appeared more subtle. We further measured their doubling times through growth curve assay in (**C**). **C**. Doubling time measurements validated the suppression interactions observed in high throughput data. The doubling times are significantly different in both suppression groups. TL variants were tested in four biological replicates, each with three or four technical replicates. The median and 95% confidence interval of each variant are shown in the figure. Kolmogorov-Smirnov test was performed to determine whether the differences in doubling time among the haplotypes showing suppression were significant. The tests resulted in a P-value of 0.0286 in both comparisons, indicating statistically significant differences. The details of growth curve experiment are in **Methods**. Specific doubling times are provided in **Supplemental Table 1**.

**Figure S9.**
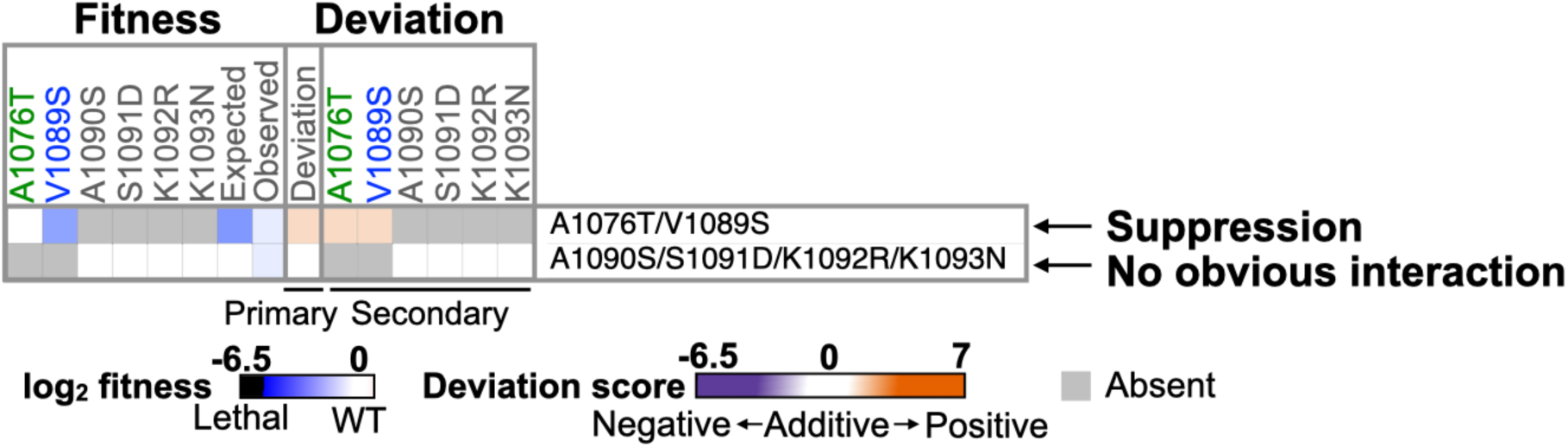
Heatmap displaying fitness and deviation scores in A1076T/V1089S and A1090S/S1091D/K1092R/K1093N. Positive deviation scores in A1076T/V1089S indicate suppression, and no deviations (primary and secondary deviation scores are around zero) indicate no interactions within A1090S/S1091D/K1092R/K1093N.

**Figure S10.**
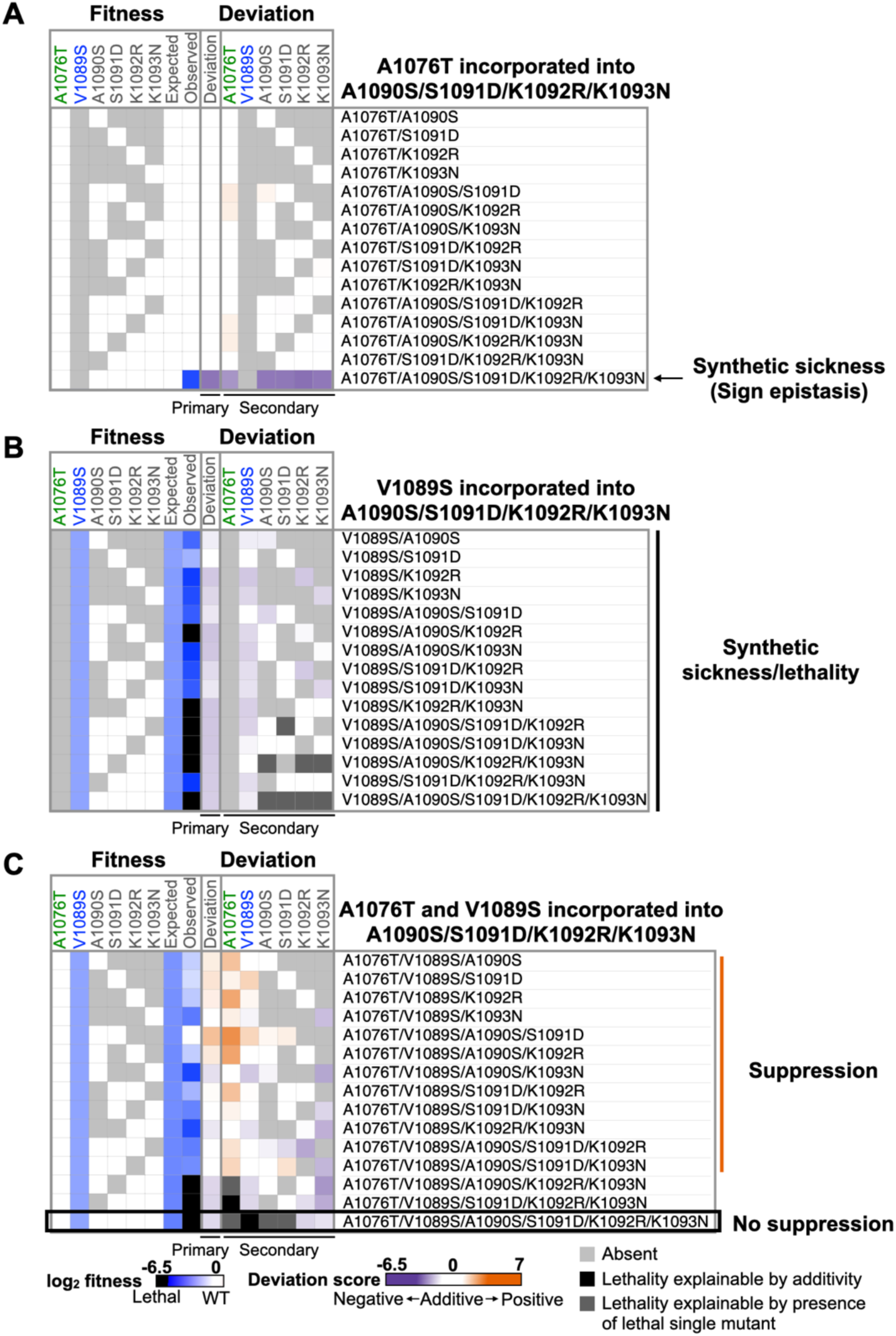
A1076T showed sign epistasis with A1090S/S1091D/K1092R/K1093N. **A**. Heatmap displays fitness and deviation scores for A1076T in all single substitutions and combinations with A1090S, S1091D, K1092R, K1093N, suggesting A1076T only showed negative interaction with A1090S/S1091D/K1092R/K1093N. **B**. Heatmap shows fitness and deviation scores for V1089S in all single substitutions and combinations with A1090S, S1091D, K1092R, K1093N. V1089S showed negative interactions with almost all of them. **C**. Heatmap shows fitness and deviation scores for A1076T and V1089S in all single substitutions and combinations with A1090S, S1091D, K1092R, K1093N. Positive interactions were observed in most combinations.

**Figure S11.**
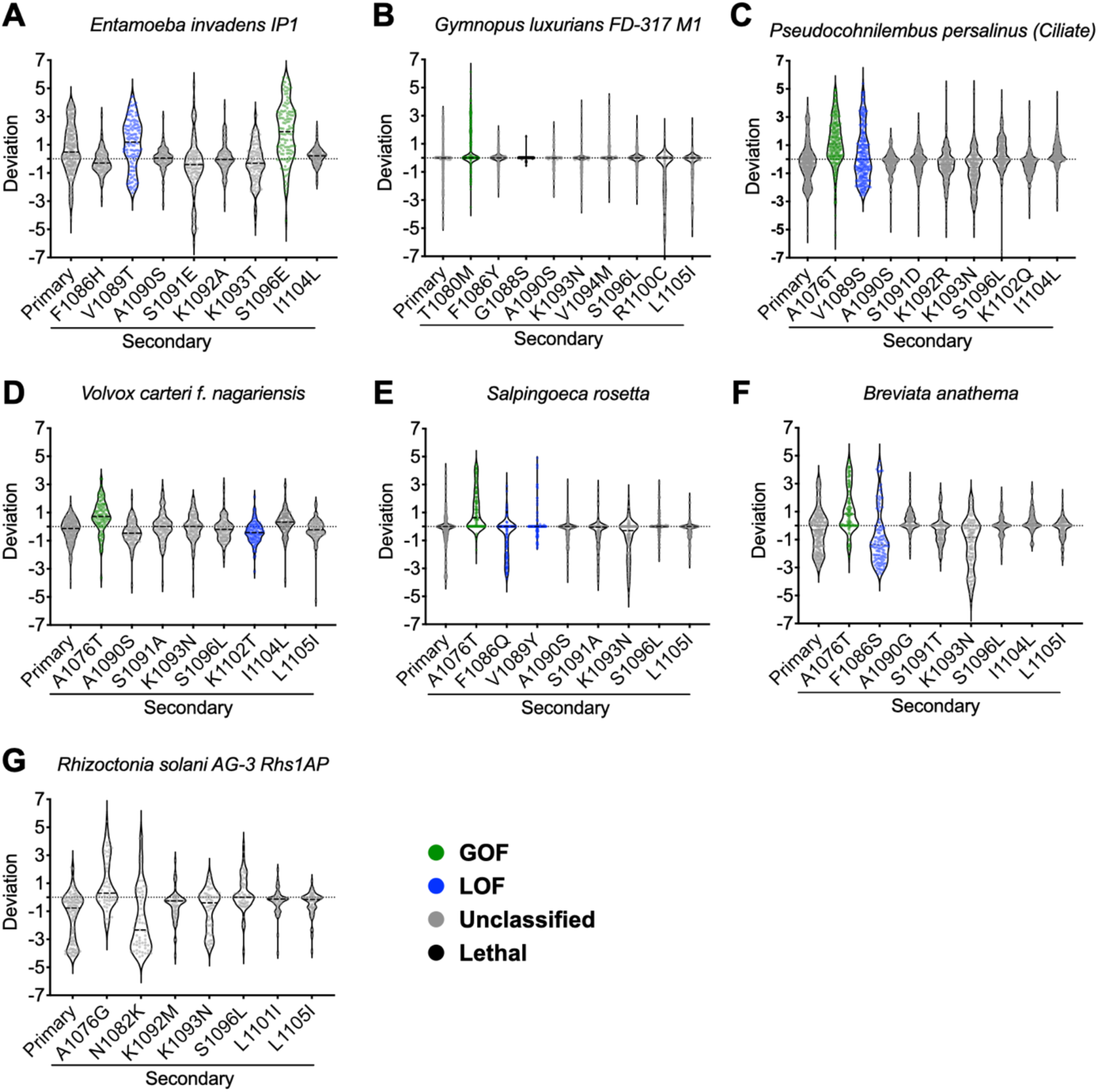
The distributions of primary and secondary deviation scores within each selected haplotype. (**A-G**). Colors of spots represent phenotypes of substitutions.

**Figure S12.**
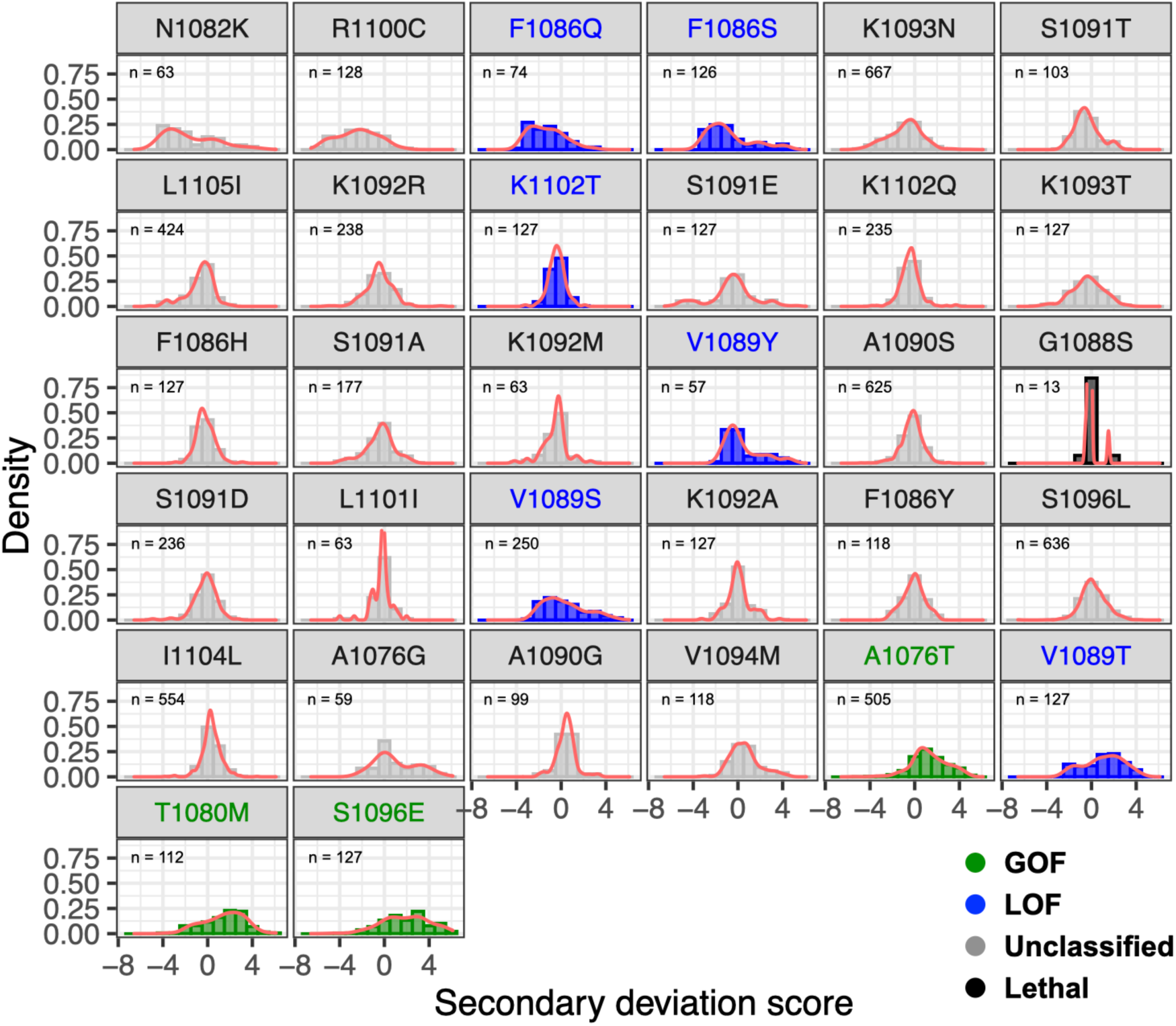
The density plots of secondary deviation scores of each substitution. We made density plots to display the distribution of secondary deviation scores for each substitution. The density plots were calculated of substitutions from nonrepetitive intermediate combinations across all haplotypes. The count of nonrepetitive intermediate combinations containing the substitution is shown in the upper left of each plot.

**Figure S13.**
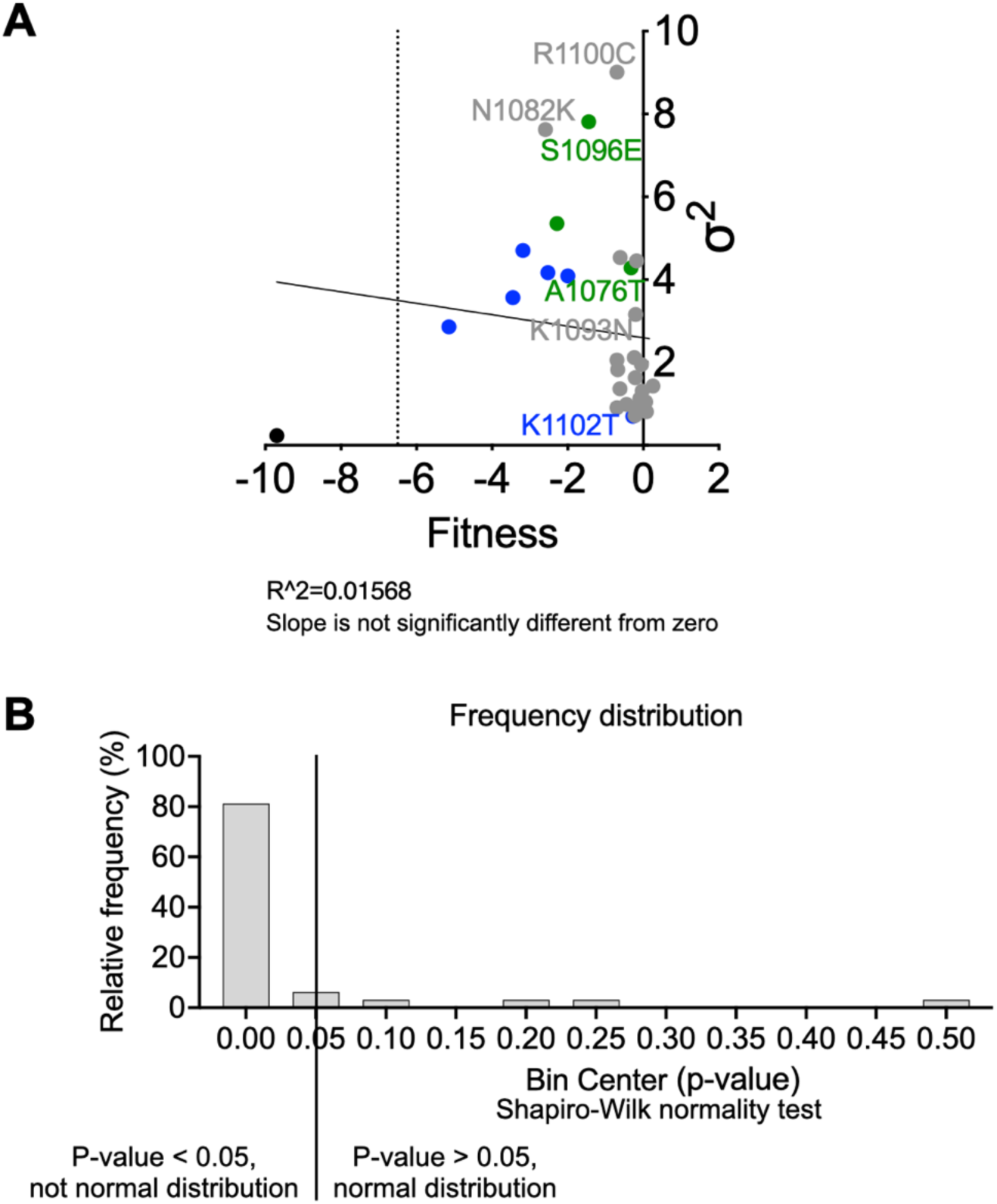
Slight but not significant negative correlation between σ^2^ and fitness of substitutions. **A**. The correlation between the maximum likelihood estimate (σ^2^) and the fitness of each substitution. Simple linear regression was applied (*y* = -0.1384\**x* + 2.600, R^2^ = 0.01568). The slope is not significantly different from zero (P = 0.5101). **B**. The histogram of p-values from normality test. The density plots of secondary deviation scores have been tested whether they follow normal distribution by Shapiro-Wilk normality test. The null hypothesis of the test is the distribution follows normal distribution. Substitutions with P-value > 0.05 suggest their distributions are normal and P-value < 0.05 indicate their distributions do not follow normal distribution. ∼80% of substitutions do not follow normal distribution.

